# High-Performance, Computer-Controlled Bipedal DNA Motor

**DOI:** 10.64898/2026.02.17.706383

**Authors:** Samrat Basak, Yaron Berger, Ofir Perel, Haggai Shapira, Toma E. Tomov, Breveruos Sheheade, Roman Tsukanov, Meitar Uralevitch, Eyal Nir

## Abstract

Achieving precise, repeatable motion at the molecular scale remains a central challenge in the development of synthetic molecular machines. Here we report a high-fidelity, fast bipedal DNA walker that moves bidirectionally along a DNA origami track using a fuel-before-antifuel operational scheme that suppresses trap states typical of externally powered DNA motors. Automated, computer-controlled microfluidics enables programmable trajectory execution with >98% yield per step and sustained bidirectional walking over distances of up to 360 nm, as monitored by ensemble FRET. Kinetic analysis reveals rapid leg lifting but slower leg placement due to inhibitory fuel–antifuel heterocomplex formation, identifying a mechanistic bottleneck that can be mitigated through optimization of fuel, antifuel and foothold design. The resulting motor operates with efficiencies of up to 4 orders of magnitude higher than previous externally controlled DNA walkers, establishing a robust framework for deterministic, programmable molecular transport.

## 1. Introduction

Biological molecular machines and motors made of proteins function in many biological processes with remarkably high chemical yield and speed.^1,2^ For example, the kinesin bipedal walker operates at a rate of hundreds of steps per second, achieving hundreds of steps before dissociating from the microtubule track.^3^ These machines and motors are autonomous, that is, they do not require external coordinating stimuli to operate properly, they are directional (that is, they overcome Brownian motion), they function as enzymes (that is, they avoid irreversible chemical change – the so called “burnt bridge” effect^4,5^), and they are typically very fast and processive. Synthetic molecular machines and devices made of various types of molecular constructs that have some of the characteristics of biological motors have been demonstrated.^6,7^ Specifically, DNA nanotechnology and DNA origami^8–14^ techniques have been utilized for rational development of programmable, complex, and functional structures, machines,^15,16^ and motors.^17–23^ Achieving a synthetic molecular machine with all the characteristics of biological machines is a very difficult task.^24^

For example, in a DNA-based approach, researchers intentionally sacrificed autonomy to achieve directionality and leg coordination while avoiding irreversible chemical changes.^14– 17,25–27^ Adapting this approach, our group has developed several generations of externally controlled non-autonomous DNA bipedal molecular walkers with the aim of achieving fast and processive bidirectional movement.^18,19,23,25,27,28^ These DNA molecular walkers operate by hybridization of the walker legs to a DNA origami-based foothold track using bridging DNA strands called ‘fuels’, followed by removal of the fuels using toehold-mediated strand displacement reaction^2^ driven by a complementary strands called ‘antifuels’.^19^

Such externally controlled DNA fuel-driven molecular motors suffer from two fundamental problems. The first is the accumulation of redundant fuel and antifuel command strands and fuel/antifuel duplex waste products in the solution; these strands lead either to motor detachment or to motors that stride in the wrong direction.^25^ We have largely resolved this issue by using a computer-controlled microfluidics device that enables removal of redundant command strands while the motor is secured on the coverslip surface,^18,29,30^ a strategy similar to a solid-phase synthesis approach.^18,28^ The automated introduction of command strands enabled by the computer-controlled microfluidics is a necessary feature when many sequential commands are required.

The second major issue is the irreversible trap-state effect. Commonly, externally controlled DNA-fuel-driven DNA motors and machines operate by first removing fuel strands using the antifuel strands, which results in lifting of a leg, followed by introduction of the next fuel strand resulting in placing of a leg; this is known as the ‘antifuel-before-fuel’ (AFBF) operational strategy.^16–19,23,25,27^ The motor can become irreversibly trapped when two fuel strands bind simultaneously to the leg and the foothold (instead of a single fuel strand that binds the leg to the foothold). A motor in such state dissociates upon the introduction of the next antifuel strand. This unwanted effect increases as fuel concentration is increased. A high fuel concentration is required for fast motor operation. Thus, the AFBF strategy results in a fundamental limitation on the performance of externally controlled DNA-fuels based motors and machines.

We have shown previously that the trap-state effect can be somewhat mitigated by using fuel strands that contain a hairpin structure^25^. These fuels first bind the foothold, then the hairpin loop opens allowing binding to the leg. Using the microfluidics delivery and the hairpin fuel approach, we designed a bipedal motor that travels on an origami track, taking a total of 32 steps, with a stepping rate of ∼6000 s and stepping yield of ∼97.5% per step, a significant performance improvement over bipedal motors operating using unstructured fuels and without washing steps.^18,25^ However, the usage of hairpin fuels did not entirely solve the trap-state problem as some hairpin fuels interact with the legs before binding the footholds, leading to some walker dissociation, especially at high hairpin fuel concentrations.^18^

Here we present a new operational scheme called fuel before antifuel (FBAF) that solves the trap-state problem entirely by reversing the order of fuel and antifuel delivery. In this scheme, the next fuel strand is added when both legs are attached to the track via the first fuel (i.e., before removal using an antifuel strand). This ensures that the introduced fuel is attached to the forward foothold but not to a leg, avoiding simultaneous attachment of two fuels and the formation of an irreversible trapped state. The excess fuels are then removed by washing using the microfluidics system, followed by introduction of the antifuel which lifts the trailing leg. This is followed by fast attachment of the lifted leg to the pre-attached forward fuel, and the walker steps forward. The FBAF strategy resulted in significantly improved speed and processivity compared to the AFBF strategy.

## 2. Results and Discussion

### 2.1. System design and implementation of the fuel-before-antifuel protocol

We designed a bipedal DNA walker system that integrates a rectangular DNA origami track with an automated, computer-controlled microfluidics platform to enable precise, repeated stepping driven by addition and removal of fuels and antifuels. The walker consists of two partially hybridized DNA strands that form a 30 base pair (bp) rigid duplex, and two single-stranded 16-base leg segments (L1 and L2). The legs are connected to the walker duplex by five-nucleotide single-stranded spacers, allowing flexibility and free rotation needed for walker stepping (**Fig. 1A**).

**Figure 1.**
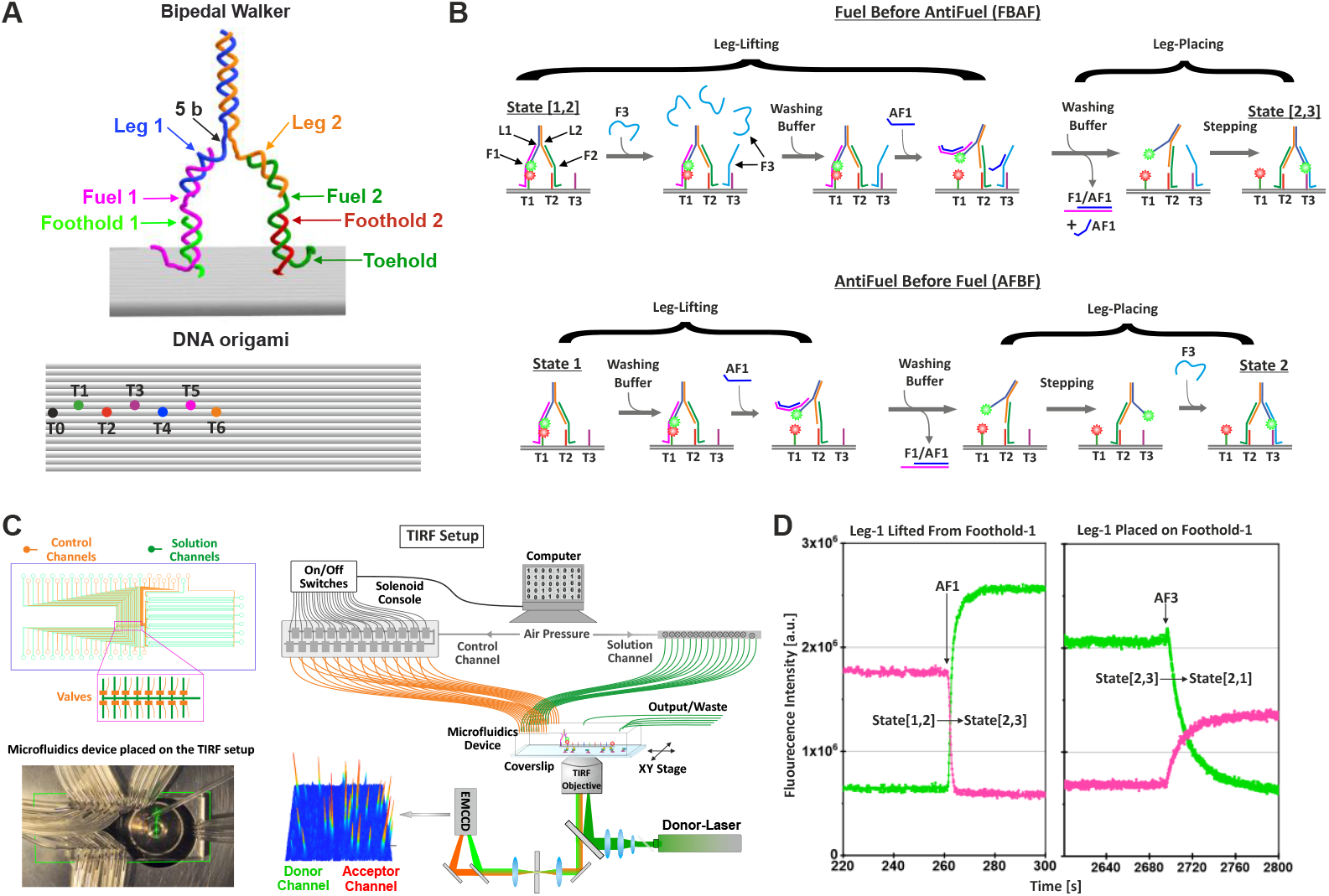
Design and operation of a DNA bipedal motor system operated with a microfluidics system and monitored using fluorescence. **(A)** Schematic of the DNA bipedal walker. The walker consists of two legs (L1, L2), each attached to footholds (T0-T6) thorough unique fuels (F1-6). The origami track includes six footholds (T1-T6) and a starting foothold (T0, for list of all DNA sequences, see **Supporting Information, Tables T1 and T2**). **(B)** Comparison of FBAF (top) and AFBF (bottom) strategies for walker operation. Schematics depict transitions from state [1,2] (i.e., walker stands on T1 and T2) to state [2,3]. **(C)** Schematic of experimental setup: The microfluidics device design (upper left), a photo of the microfluidics device mounted on the microscope objective (bottom left), schematic of the entire experimental system, including the optical setup (right). **(D)** Donor (green) and acceptor (magenta) fluorescence time traces (signals are summed over ∼100 motors) for motor operated using the FBAF strategy. Lifting of donor-labeled L1 from acceptor-labeled F1 (i.e., walking forward from state [1,2] to state [2,3], left panel) and placement of L1 on F1 (i.e., walking backwards from state [2,3] to state [1,2], right panel) are shown.

The rectangle origami track measures 40 × 150 nm and contains a starting foothold (T0) and six unique foothold strands (T1–T6), each separated by 12 nm, a distance previously optimized for best performance.^28^ This origami was designed such that it can include six additional footholds (for a total of 13 footholds) to cover the entire origami length, but only one set of six footholds was used in this work (**Supporting Information, Figure S1**). Footholds are created by extending selected origami staples with sequences containing segments complementary to the matching fuels, enabling foothold–fuel–leg binding.

In the FBAF strategy (**Fig. 1B, top**), fuel strand F3 is introduced while both walker legs are bound to the track (L1 to T1 and L2 to T2), allowing F3 to hybridize with the unoccupied forward foothold T3. After this pre-loading step and removal of the excess F3, the corresponding antifuel strand, AF1, is introduced. This displaces the fuel F1 from the trailing leg L1 and allows this free leg to immediately bind to the preloaded fuel F3. This operation sequence avoids the trap-state scenario, prevalent in the more common AFBF scheme (**Fig. 1B, bottom**), where two different F3 strands might bind the leg L1 and the foothold T3, blocking the desired leg-foothold interaction, a process that leads to walker dissociation from the track in the following step (**Supporting Information, Figure S2**). Note that the antifuel strands are 8 bases shorter than the corresponding fuel strands, which accelerates the leg-placing reaction as discussed in section 2.4.

To implement the complex timing and sequence control required, we used a custom pneumatic microfluidics device with 33 independently controlled input channels. Each channel is linked to a reagent reservoir via a computer-controlled solenoid valve, allowing rapid switching between fuels, antifuels, washing buffers, and the ingredients for immobilizing the motor inside the microfluidics device (**Supporting Information, Figure S3 and Table T3**). By automating the delivery sequence (fuel → wash → antifuel → wash) and controlling incubation and washing times precisely, the system ensures high reproducibility and minimizes cross-contamination (**Fig. 1C**).

Ensemble Förster resonance energy transfer (FRET) total internal reflection fluorescence (TIRF) readout was used to monitor walker stepping in real time. L1 and T1 were labeled with donor and acceptor fluorophores, respectively, such that high FRET signal indicates L1 positioned on T1 and low FRET signals indicates L1 is not bound to T1. The progress of the motor is monitored by measuring the fluorescence intensities of the donor and acceptor fluorophores of ∼100 individual motors immobilized inside the microfluidics device (**Fig. 1D**). The lifting of L1 from T1 and placing of L1 on T1 can be clearly detected, and the reaction profiles are measured in real time. In cases of complete dissociation of a walker from the track (i.e., an operational error), the donor and acceptor signals are reduced, allowing measurement of stepping yields.

### 2.2. High-fidelity, reversible stepping under fuel-before-antifuel operational strategy

To assess the performance of our motor under the FBAF operation scheme, we commanded the walker to walk from state [1,2] to state [4,5] and back to state [1,2] five times (**Fig. 2A**; for the procedure used for the initial positioning of the walker at state [1,2] see the **Supporting Information section 1.9 and Figure S4**). At state [1,2], the FRET signal was high, and in all other states the FRET signal was low, as expected from the positions of the fluorophores in these states (**Fig. 2B**). There was a dip in donor intensity in the middle of the time trace that was not related to the placing or lifting of L1 from T1. We attributed this decrease in donor intensity to photophysical effects related to the positions of the fluorophores in respect to the DNA construct (see detailed explanation in **Supporting Information section 1.10 and Figure S5**).

**Figure 2.**
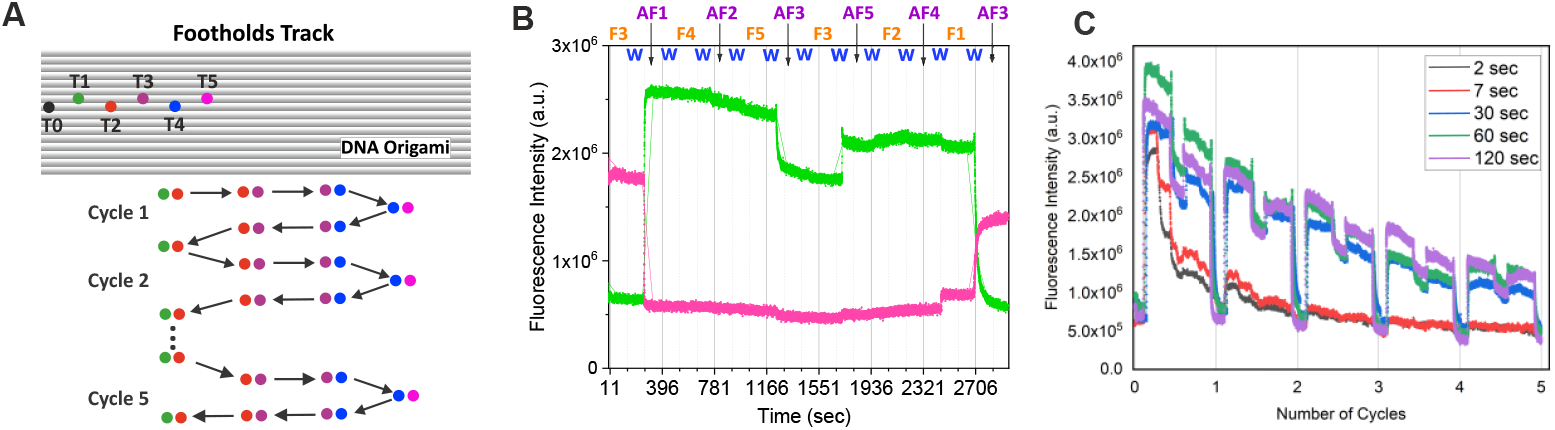
Analysis of the bipedal walker system performance. **(A)** Schematic of the back-and-forth movement of the bipedal walker from T1 to T5 (for full list of microfluidics commands, see **Supporting Information, Table T4**). **(B)** Donor (green) and acceptor (magenta) fluorescence time traces. **(C)** Donor fluorescence intensity profiles measured over five walking cycles at fuel and antifuel incubation times of 120, 60, 30, 7, and 2 s.

The motor operation was measured under different fuel and antifuel incubation duration times and washing duration times (120, 60, 30, 7 and 2 s), and the survivability of the walker on the track (i.e., stepping yields) were estimated by measuring the relative donor intensities upon L1 placing and lifting (**Fig. 2C**; for the method used to calculate stepping yields, see **Supporting Information section 1.11 and Figure S6**).

For longer incubation times (120, 60, 30 s), the survival rates were high enough to allow at least five back-and-forth walking cycles that included 30 steps (i.e., 60 leg-lifting and leg-placing reactions). This amounts to a total walking distance of 360 nanometers. For the 120-s incubation time, the calculated average stepping yield was ∼98%, which means that only ∼2% of the walkers dissociated at each step. At the incubation time of 7 seconds, the walker dissociation was ∼10% per step, and at the 2-s incubation time, the dissociation rate was around 25%. Importantly, the data clearly showed that the fuel addition steps did not lead to any significant changes in the fluorescence intensities, confirming that the leg remained stably bound until an antifuel-mediated leg release was triggered.

### 2.3. Kinetic analysis reveals rate asymmetry

Although the FBAF walker achieved high stepping yields under generous incubation times, at shorter incubation times stepping yields dropped sharply. To understand this operational bottleneck, we measured and analyzed the leg-lifting and the leg-placing reactions rates separately.

To measure the leg-lifting reaction profile, the walker was positioned at a high-FRET state [1,2]. At time zero, we introduced 0.1, 0.3, 1, 5, or 10 µM antifuel AF1; this resulted in a drop in FRET signal, indicating that L1 was lifted from T1 (**Fig. 3A, upper panel**). To calculate the observed reaction rates, the FRET kinetic profiles were fitted to single-exponential functions, and the resulting rates were plotted against the AF1 concentrations to calculate the concentration-independent leg-lifting reaction rate (**Fig. 3A, middle and bottom panels**). The rate, k_LL_, was 0.34 ± 0.09 × 10^6^ M^-1^ s^-1^. This value agrees with our previous measurements of the leg-lifting reaction rate in AFBF system^18^ and is typical for DNA strand hybridization and toehold-mediated strand displacement.^17,24,25^ The single-exponential characteristic of the reaction and the linear dependence of the leg-lifting reaction rates on the AF1 concentrations indicate that the leg-lifting process was fully completed within a time frame consistent with a first-order dissociation mechanism, with rates depending only on the binding rate of the antifuel to the foothold, in agreement with our previous finding.^18,25^ In other words, the antifuel-mediated fuel removal reaction is very efficient and fast, and it does not constitute a major limitation on motor speed or yield.

**Figure 3.**
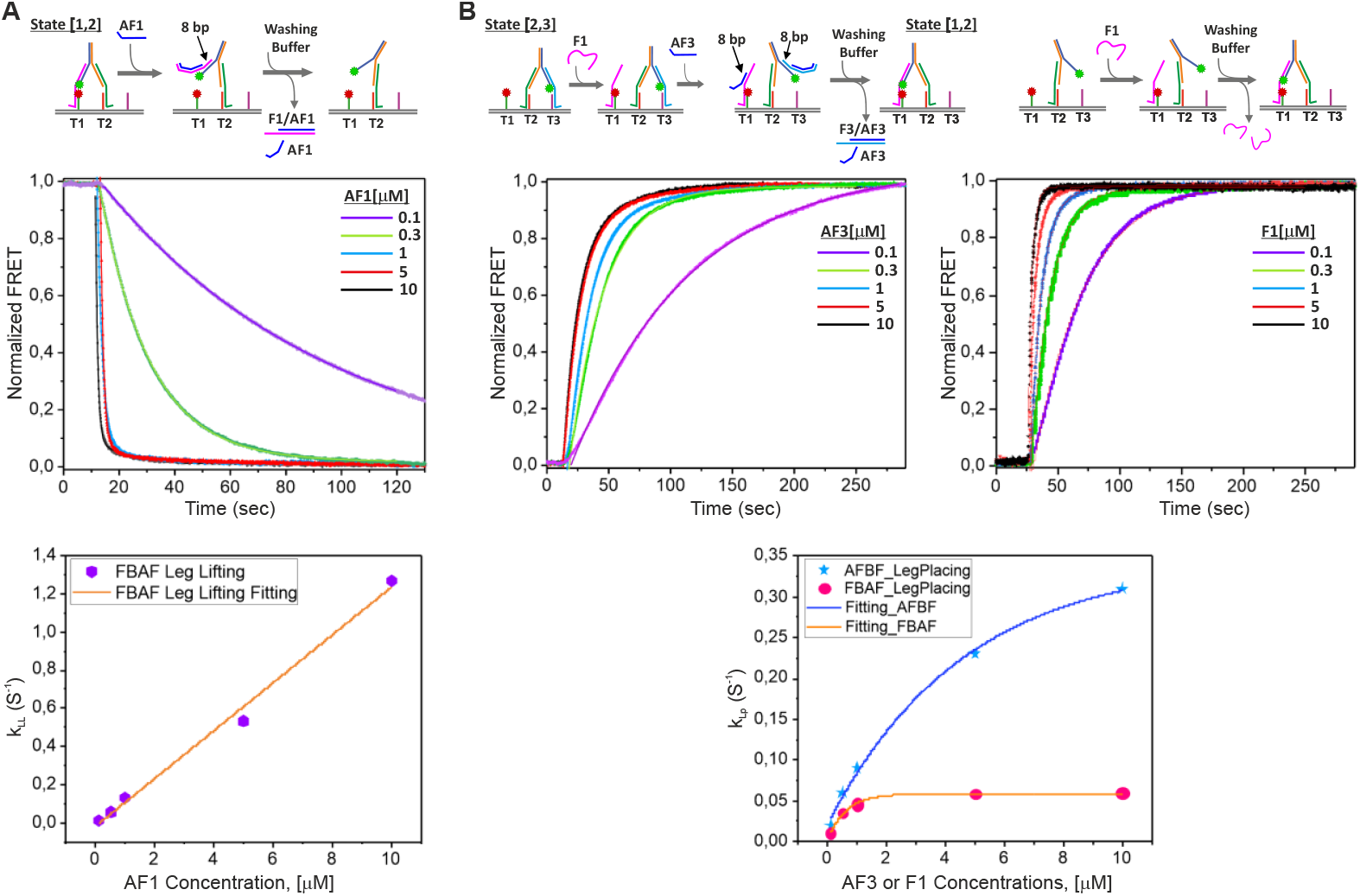
Leg placing and leg lifting kinetics. **(A)** Schematic of kinetic measurement of FBAF leg-lifting rate (upper panel), FRET kinetic profiles of leg-lifting reactions over a range of AF3 concentrations fit using an exponential function (middle panel), and summary of the calculated rates (bottom panel). **(B)** Schematic of kinetic measurements of FBAF and AFBF leg-placing reactions (upper left and right panels, respectively), FRET kinetic profiles over a range of concentrations of AF3 or F1 (middle left and middle right panels, respectively), and summary of the calculated rates (bottom panel).

In the FBAF scheme, it is impossible to measure the leg-placing reaction alone because the fuel must be introduced while both legs are still attached to footholds, and stepping is initiated by antifuels. Therefore, the leg-placing reaction rate can be measured only together with the leg-lifting rate (which is, fortunately, significantly faster) in the context of a full stepping reaction. To measure the rate, the walker was positioned in a low-FRET state [2,3], and F1, which binds to T1, was added followed by washing to remove excess fuel strands. To initiate the stepping reaction, AF3 was introduced at time zero. This resulted in rapid lifting of L1 from T3, followed by the slower leg placing reaction (i.e., L1 placed on T1 by binding F1), achieving state [1,2]. Experiments were performed at 0.1, 0.3, 1, 5, and 10 µM concentrations of AF3 strands (**Fig. 3B, upper and middle left panels**). The kinetic profiles were fitted to exponential functions, and the observed rates were plotted against the AF3 concentrations (**Fig. 3B, bottom panel**). The stepping reaction rates were significantly slower than the leg-lifting rates, indicating leg-placing are significantly slower than leg-lifting reactions. The stepping rates increased with increased AF3 concentrations, but only to a certain asymptotic value (k_lp_ [maximum] = 0.06 s^-1^, equivalent to 16.7 ± 4.2 s minimum reaction time), indicating that the rate-limiting step was the leg-placing reaction.

For comparison, we measured the leg-placing reaction rates for the AFBF system. The experiment began with the walker standing on L2 with L1 lifted. At time zero, F1 was introduced, resulting in binding of L1 to T1 (**Fig. 3B, upper and middle right panel**). Because of the formation of trapped states, an increased concentration of F1 resulted in decreased yield (as seen in the non-normalized FRET curves **Supporting Information, Figure S2**); however, the leg-placing reaction rates were noticeably faster than rates in the FBAF system (**Fig. 3B, bottom panel**).

### 2.4. Formation of unwanted fuel/antifuel heterocomplex

Although the FBAF strategy is superior to the AFBF strategy, FBAF involves an unwanted side reaction that limits its performance. When antifuel (i.e., AF3) is introduced, it not only binds fuel F3, but may also bind the pre-positioned backward fuel (i.e., F1), forming an unwanted AF3/F1 heterocomplex (HC1, **Fig. 4**). This interaction is unavoidable because F1 and F3 contain an identical segment complementary to L1, and, therefore, AF3 is not only complementary to F3 but also complementary to the exposed part of F1. Thus, antifuel AF3 not only removes fuel F3 from the system but may also block the pre-positioned fuel F1 from binding the leg L1.

**Figure 4.**
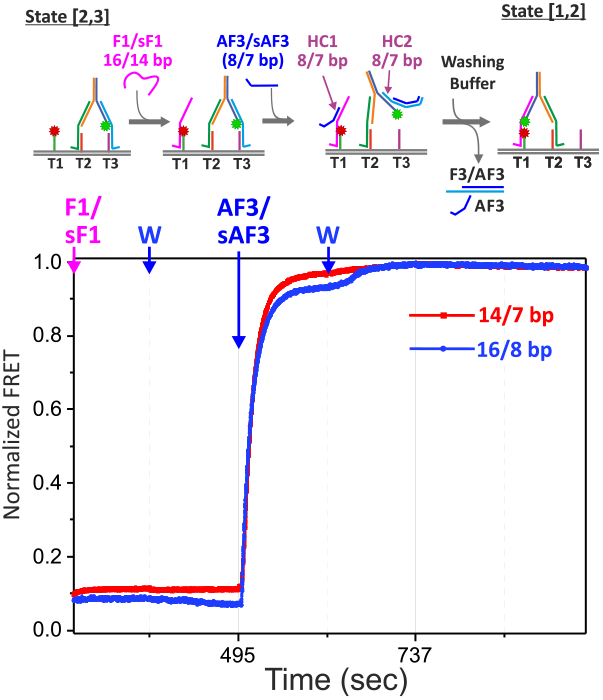
Heterocomplexes inhibit motor operation. Schematic of the motor system and the heterocomplexes formed during FBAF operation (upper panel). Normalized FRET profiles of leg placing reaction (L1 on F1) measured for longer fuels and antifuel (16/8 system, blue) and shorter fuels and antifuel (14/7 system, red curve).

To mitigate formation of this unwanted heterocomplex, FBAF antifuel strands were designed to be shorter than the corresponding fuel strands by 8 bases, resulting in a heterocomplex of 8 bp (instead of 16 bp for a full length antifuel strands). The shorter duplex accelerates the leg-placing reaction by two ways: First, it destabilizes the heterocomplex, promoting thermal dissociation of the antifuel from the fuel. Second, the unbounded 8-base segment of the fuel F1 is free to interact with L1 even before the dissociation of the heterocomplex. Shortening the antifuel strands by 8 bases also results in some incomplete removal of the fuel from the leg, but this 8-bp fuel/leg heterocomplex (HC2, **Fig. 4**) readily thermally dissociates, freeing the leg L1 to interact with the backward fuel F1. A careful observation of the FRET kinetic profile of the leg-placing reaction shows that a small percentage of walkers do not complete the leg-placing reaction before the excess antifuel strands (AF3) are removed; it is only after washing that L1 binds fuel F1 (**Fig. 4**). These data support our hypothesis that there is transient formation of an unwanted heterocomplex-1.

To further test this hypothesis, we measured walking driven by shorter fuels, sF1 and sF3, which were design to bind L1 through 14 bp (instead of 16 bp in the F1/L1 and F3/L1 complexes), and a shorter antifuel strand sAF3, which was designed to remove 7 bases from the F3/L1 complex (instead of 8 bases with AF3). With this design both heterocomplexes were of 7 bp in length, which should accelerate heterocomplex dissociation relative to the longer fuel and antifuel strands. The results clearly show that with these shorter strands, the leg-placing reaction reached completion faster than with the longer strands (**Fig. 4**).

### 2.5. Solution for the heterocomplex problem

The dissociation rate of a duplex DNA is exponentially dependent on its thermodynamic stability, and, therefore, its length in base pairs. For short duplexes, a reduction in length by one base pair typically increases the dissociation rate by a factor of 3 to 5 (dependent on whether the base pair removed is G:C or A:T). Reduction in the length of the heterocomplex duplex by one base-pair (from 8 to 7 bp) may not destabilize the heterocomplex enough to completely prevent interference with the stepping reaction, but reducing the duplex length to 6 bps may weaken the leg/fuel interaction enough that the legs may thermally dissociate from the footholds, resulting in walker dissociation.

To solve the heterocomplex problem, we propose the fuels before 2 antifuels (FB2AF) strategy. In the FB2AF strategy, the fuel strands are designed to bind the legs through 15 bps (instead of the 18 and 16 bp used previously) and contain two toehold segments, one on each end of the fuel strands (**Fig. 5**). Two corresponding antifuel strands (i.e., AF1a and AF1b) would bind these two toeholds, leaving only 5 bps F1/L1 duplex. This duplex, and the 5 bps AF1a/F3 and AF1b/F3 duplexes, are expected to thermally dissociate at in less than 1 s^31–34^, potentially solving the heterocomplex problem.

**Figure 5.**
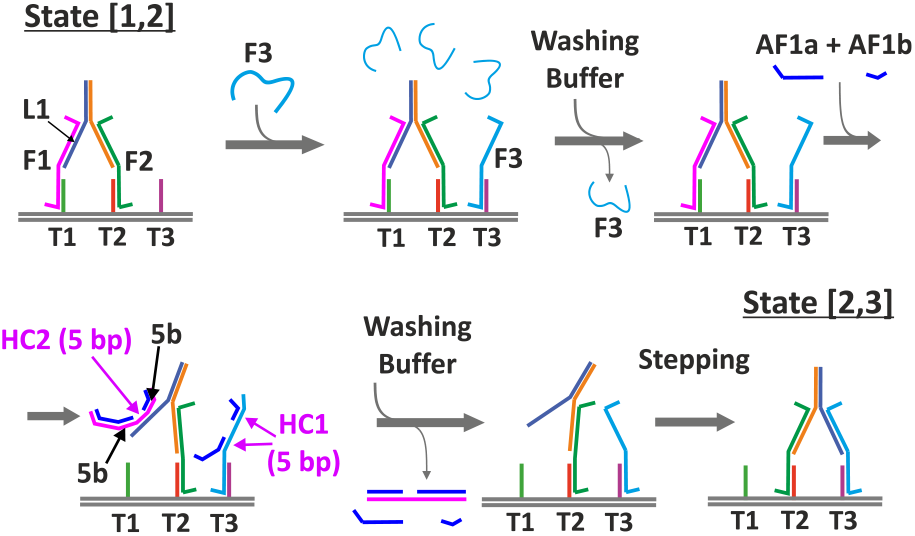
A schematic of the proposed FB2AF mechanism. The fuel strands contain two foothold segments, one at each end of the fuel. Two antifuel strands, AF1a and AF1b, are introduced that together removing the fuel F1 from both leg L1 and foothold T1. With this design, heterocomplexes (HC1, HC2) are only 5 bps and are predicted to thermally dissociate exponentially faster than the heterocomplexes formed in the FBAF approach.

### 2.6. FBAF versus AFBF system performance

To improve performance of the DNA motor, we focused on speed and processivity (i.e., walking yield). The performance of the FBAF motor developed in this work was compared to that of our previously described AFBF motor.^18,25^ For the AFBF system, high fuel concentrations allow fast walking, but because of the trap-state effect, this comes at the expense of a high percentage of walkers dissociating from the track at each step (∼20% at 10 µM fuel concentration, **Fig. 6, green curve**). Lowering the fuel concentration improved walking yields at long incubation times, but yields dropped significantly at short incubation time, limiting motor speed (**Fig. 6, black curve**). Unlike the AFBF motors, the FBAF motors does not form the trapped state reducing tradeoff between speed and yield (**Fig. 6, red curve**). The FBAF system had between 0.5 and 1.5 orders of magnitude performance improvement over the AFBF motors, depending on the concentration of fuel used to operate the AFBF motor. For the 100 s per step walking rate, the FBAF shows ∼5% walker dissociation per step, whereas the AFBF mechanism shows ∼20%. For the AFBF motor to achieve 5% dissociation rate per step, the fuel concentration must be 0.2 µM (or lower) and the stepping rate must be reduced to a step per ∼3000 s (or slower, **Fig. 6, black curve**).

**Figure 6.**
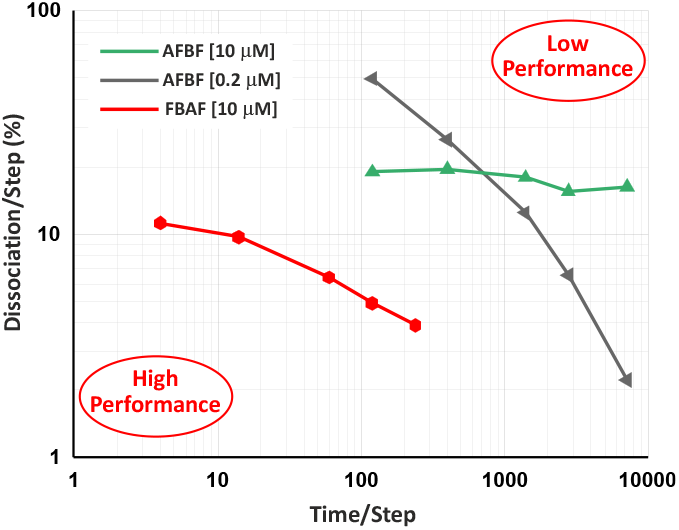
Comparison of the performances of the FBAF and AFBF motor systems. Shown are the walker dissociation rates per step measured for motors to operate at different stepping rates. The bottom left corner represents high performing motors, and the top right corner represents low performing motors. The AFBF system was tested at 0.2 and 10 µM fuel and antifuel concentrations (black and green curves, respectively), and the FBAF system was tested at 10 µM fuel and antifuel concentrations (red curve). The AFBF data is taken from Tomov et al.^18^

## 3. Conclusions

We have previously shown that AFBF motors perform much better when operated using a microfluidics device that allows the removal of excess fuels, antifuels, and fuel/antifuel duplex waste.^18^ The trap-state effect limited the performance of this motor system, however. The high fuel concentration necessary for a rapid stepping rate also resulted in low stepping yields. The FBAF system allows use of a high concentration of fuels without any reduction in stepping yields, significantly reducing coupling between speed and processivity, and resulting in 0.5-1.5 orders of magnitude performance improvement over previous AFBF motors. The formation of unwanted heterocomplexes in the FBAF system places a new limitation on motor performance. This limitation can be mitigated to some extent by reducing the length of the fuel/antifuel and fuel/leg interacting segments such that the heterocomplexes are destabilized. We suggest that using two antifuel strands that interact simultaneously with the fuel (the FB2AF system) will result in further destabilization of the heterocomplexes, increasing motor performance even further. We have recently developed a DNA rotor made of two disk-like DNA origami that uses two sets of bipedal walkers that operate using the FBAF mechanism developed in this work. The rotor operation was very robust. For example, it could perform eight full rotations, amounting to 96 steps, with a natural ability to recover from operational error,^23^ demonstrating the advantages of using the microfluidics and the FBAF approach. Our group has been working to understand and improve externally controlled DNA bipedal walkers for more than a decade.^18,19,23,25,27,31^ As demonstrated here, we have improved motor performance by 3-4 orders of magnitude relative to the motors that operate without microfluidics that employ the conventional AFBF approach and non-hairpin fuels.^25^ We are confident that the FB2AF system will improve walker performance even further and that a system with less than 1% error rate per step at a stepping rate of several seconds per step is within reach.

## Acknowledgements

The authors gratefully acknowledge financial support from Israel Science Foundations (ISF), through the “High-Performance Linear and Rotary DNA Nanomotors” project. We thank Doron Gerber and Noa Marchoom-Bura from Bar-Ilan University for providing microfluidic devices.

## Data availability

Data used for the research is available upon request.

## Additional information

Supplementary information is available for this paper.

## Notes

### Competing Interest Statement

The authors have declared no competing interest.

### Summary of Updates

The work is done by ensemble FRET, and in the abstract it is written as single-molecule FRET, so the necessary changes has been made, and the manuscript is updated to the current version.

