## Supplementary Material for "High-Performance, Computer-Controlled Bipedal DNA Motor"

#### 1.1. DNA synthesis and labeling

PAGE-purified DNA components, including footholds, legs, fuels, antifuels, and biotinylated origami anchor strands, were purchased from Integrated DNA Technologies (IDT). Specific DNA strands (L1, T1, T7, and T13) were synthesized with a C6dT internal amino modifier (iAmMC6T) at desired positions and were purified using HPLC by IDT. For fluorescent labeling, the amine-modified DNA strands were dissolved in 100 mM sodium tetraborate at pH 8.4 and mixed with a DMSO solution of either ATTO-550 or ATTO-647N dye (donor or acceptor, respectively; ATTO-TECH). After incubation overnight at 70 °C in the dark to facilitate the conjugation reaction, the DNA-dye conjugates were precipitated using 100% methanol. The labeled DNA pellet was isolated from the excess dye by centrifugation, then resuspended in 100 mM TEAA with 5% acetonitrile and purified using a reverse-phase HPLC column (Xterra C18, Waters). The labeling efficiency was typically around 85%, and the purity of the labeled products was greater than 99%, as confirmed by reinjection into the HPLC system. The purified labeled DNA samples were lyophilized, resuspended in 10 mM Tris-HCl, 1 mM EDTA, pH 8.0 (TE buffer), and stored at -20 °C for further use.

### 1.2. DNA origami design

The previous iterations of our DNA motor utilized Rothemund's rectangular origami design, measuring 60 × 90 nm[1]. However, this design had certain limitations due to its inherent right-handed global twist, which arises because the design twist density is set at 10.66 base pairs per turn, slightly deviating from the natural B-DNA twist density of 10.5 base pairs per turn. This discrepancy led to difficulties in the assembly of extended tracks from multiple origami tiles. To correct the global origami twist and achieve a twist density of 10.5 base pairs per turn, a base deletion strategy was employed. Specifically, one base was removed every 64 bases (or 1 base per 8 base blocks), effectively aligning the twist density with that of B-DNA. Additionally, we previously showed that foothold spacing of 12 nm allow the fastest leg placing and, therefore, most efficient motors[2]. To prevent the walker from skipping between foothold sets, a minimum distance of 45 nm between identical footholds in consecutive sets is required. To accommodate these parameters and ensure that the origami construct can include at least two complete 6-foothold sets (for future studies), we designed the origami to be of a 40 × 150 nm size (**Figure S1A**). The new structure was assembled using M13mp18 single-stranded DNA (ssDNA) as the scaffold (New England BioLabs), with non-purified staple strands obtained from IDT. The structural design of the DNA origami was performed using the computer-aided design software CaDNAno[3]. To verify the accuracy of the predicted three-dimensional configurations, the designs were further analyzed using MrDNA[4], a multi-resolution model that predicts the structure and dynamics of DNA constructs (**Figure S1B** ).

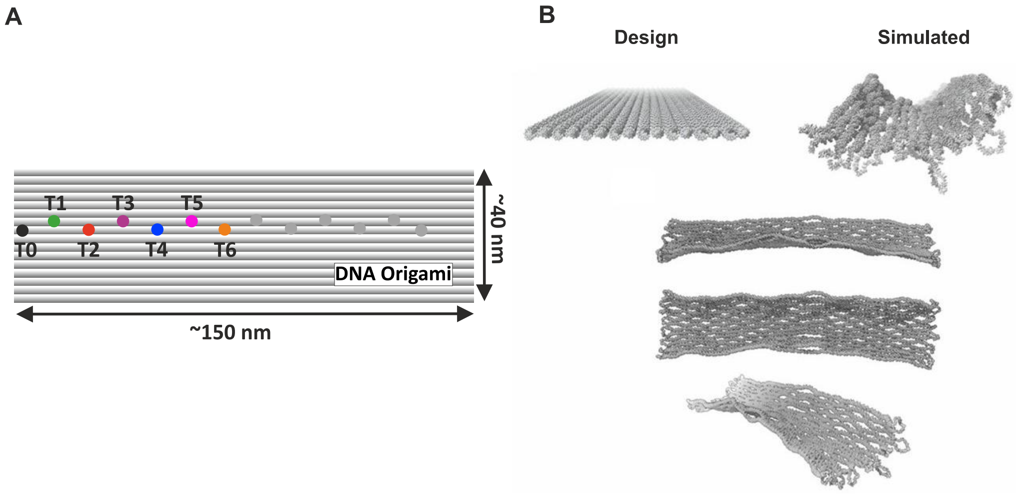

**Figure S1.** Design and simulation of the origami construct. **(A)** Schematic of DNA origami tracks. Only the first set of 6 footholds (T1-T6) was used in the experiments described here. The second set is grey. **(B)** Comparison of idealized design of the origami structure and DNA origami structures predicted by the mrDNA coarse-grained simulation framework. The top left panel shows the designed rectangular origami architecture, while the other panels illustrate the simulated conformational state view from different angles.

### 1.3. Optimized annealing conditions for DNA motor assembly

The annealing solution was prepared with a final composition of 2 nM scaffold DNA, a 5-fold molar excess of staple strands, a 10-fold molar excess of foothold strands, a 16-fold molar excess of walker leg strands, and a 20-fold molar excess of fuel strands. The solution was prepared in a buffer containing 40 mM Tris, 1 mM EDTA, and 40 mM acetic acid, pH 8.0 (1X TAE) supplemented with 12.5 mM $MgCl_2$, in a final volume of 50 µL (5). The annealing procedure was carried out in a PCR machine under the following conditions: an initial incubation at 65 °C for 15 min, followed by a gradual cooling to 40 °C at a rate of 1 °C per 15 min, and further cooling to 20 °C at a rate of 1 °C per 10 min.

### 1.4. Post-annealing purification of DNA origami structures

Following annealing, the DNA origami structures were purified using a polyethylene glycol (PEG) precipitation method to remove excess staples, footholds, fuel strands, and walker components, which could otherwise interfere with the proper functioning of the motor. After annealing, the solution was mixed 1:1 (v/v) with a precipitation buffer consisting of 5 mM Tris, 1 mM EDTA, and 500 mM NaCl, pH 8.0 supplemented with 15% PEG 8000 (w/v; Sigma-Aldrich). The mixture was then centrifuged at 17,800 g for 30 minutes at 4 °C in a microcentrifuge (Eppendorf 5417R). The supernatant was carefully removed, and the DNA origami pellet was resuspended in a buffer containing 1X TAE and 15% PEG in a 1:1 ratio for an additional round of purification to ensure thorough removal of contaminants. The final DNA origami pellet was resuspended in 10 mM Tris, 1 mM EDTA, 3 mM Trolox, and 100 mM NaCl, pH 8.0 (working buffer) and incubated in a shaker incubator at 25 °C for approximately 30 min to achieve complete dissolution.

### 1.5. Design and functionality of the microfluidics device for DNA motor operation

The microfluidics device used in this study was originally developed in collaboration with Dr. Doron Gerber from Bar-Ilan University.[18] Initial versions of the device were limited by the number of channels,[5,6] which are insufficient for operating a motor with six distinct footholds. To address this limitation, a device was constructed with six working channels (instead of the four in previous work[6]) and additional input lines, enabling a broader range of experiments to be conducted on the same chip. The working channels were arranged in a "zigzag" pattern to maximize the surface area available for measurement. The microfluidics device is fabricated from silicone elastomer polydimethylsiloxane and consists of two layers: a bottom layer containing the solution channels and a control layer with integrated pneumatic valves. The input channels deliver reagents necessary for immobilizing, assembling, and operating the DNA motor, including washing buffers, BSA, NeutrAvidin, DNA origami tracks, walker strands, fuel strands, and antifuel strands. The input channels and working chambers are interconnected via a feed channel that includes a drain outlet. Each input channel and the connections to and from each working chamber are regulated by pneumatic Quake-type valves. The pneumatic valves are operated by varying the back pressure. Air pressure is controlled by solenoids located outside the microfluidic device. When pressure is applied, the valve expands to obstruct the flow of solution through the channel. Conversely, when a solenoid is closed, the valve opens, allowing free flow of the solution in the designated channel. The volume and flow rate of solutions are precisely controlled by adjusting the back pressure and flow duration, allowing for the controlled assembly and operation of the motor as well as the sequential introduction and removal of an essentially unlimited number of solutions (e.g., fuels and antifuels).

#### 1.6. FRET-TIRF experimental setup and synchronization

The FRET-TIRF experiments were conducted using a custom-built optical setup.[6–9] A continuous-wave green laser beam (532 nm, MLL-FN-532, Changchun New Industries Optoelectronics Tech. Co., Ltd.) and a red diode laser (640 nm, Cube 640-40C, Coherent Europe) were aligned into a single-mode fiber (Thorlabs). The combined beam was then collimated and expanded by a factor of 4.16 before being focused off-axis through an achromatic lens (180 mm, AC508-180-A, Thorlabs) onto the back aperture of a high numerical aperture oil immersion objective (NA 1.45, 100×, Olympus America) mounted on a commercial inverted microscope (IX71, Olympus America). During measurements, the laser intensity was set to 25 μW to minimize photobleaching effects. The fluorescence emitted from the sample was collected and separated from the excitation light using a dichroic mirror (ZT532/638RPC, Chroma). The collected light was then filtered (ZET 532/642 M, Chroma) and split into donor and acceptor wavelengths using a second dichroic mirror (645 DCXR, Chroma). The donor and acceptor fluorescence signals were further refined by band-pass filters (580/60 and 731/137, respectively; Semrock) and focused onto an electron-multiplying charge-coupled device camera (IXON DU-897E, Andor). The donor channel was aligned to the left side of the camera chip, and the acceptor channel was aligned to the right. The microfluidic device was securely mounted onto the microscope stage (IX-SVL2, Olympus) using slide clips (IX-SCL, Olympus). This setup allowed for precise XY translation of the microfluidics relative to the field of view, facilitating imaging at various locations along the working chamber. The optical setup included two custom-built shutters controlled by a LabView-based microfluidics valve control program. These shutters allowed independent or combined operation of the green and red lasers at different stages of the motor operation, effectively reducing photobleaching during prolonged data acquisition (approximately 10,000 s). To synchronize data collection, the Andor camera was set to begin recording .TIFF files upon receiving the first input signal from the microfluidics control valves. This synchronization was achieved using the "External Start" option on the camera, with triggering controlled by a 5-volt transistor-transistor logic signal (TTL) generated by an Arduino (UNO Rev. 3). This precise timing coordination ensured that the recorded fluorescence spectra were accurately aligned with the input commands from the microfluidic system, enhancing the reliability and reproducibility of the experimental measurements.

### 1.7. The trap-state effect

The trap-state mechanism is shown in **Figure S2A**. When the fuel strand F3 is introduced, it binds either the complementary leg L1 or the complementary foothold T3, and then, the same fuel strand binds the foothold T3 or the leg L1, completing the stepping reaction. However, a second strand of the fuel may bind to either the leg or foothold, preventing leg placing and leading to a trapped state. In such a case, the walker dissociates from the track upon the introduction of the consecutive antifuel strand AF1. An increase in the concentration of fuel F3 results in a faster leg placing reaction but at the expense of lower yields (**Figure S2B**).

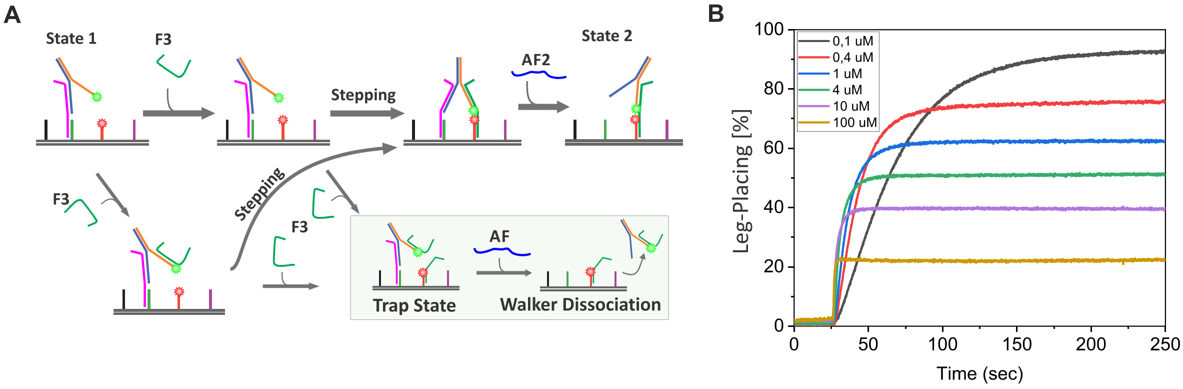

**Figure S2.** The trap-state effect. **(A)** Schematic of formation of the trapped state and subsequent walker dissociation. **(B)** FRET kinetic profiles of leg placing reaction (L1 on T3) measured at different fuel (F3) concentrations.

#### 1.8. Immobilization of DNA origami motors in the microfluidic device

The immobilization of DNA origami motors onto the microfluidic device was achieved through non-covalent avidin-biotin interactions (**Figure S3**). The DNA origami structures were modified with biotinylated anchor strands on the bottom surface, opposite the foothold surface, to facilitate this attachment. The immobilization process involves several carefully controlled steps. First, the microfluidic working channel was hydrated with 10 mM Tris, 50 mM NaCl, pH 8.0 (T50 buffer) to prepare the surface. Following hydration, a solution of biotinylated BSA (1 mg/mL, A8549, Sigma-Aldrich) was flowed through the channel for 15 min resulting in a surface covered with biotin. In some experiments, a 1 mg/mL solution of biotinylated PEG (Biotin-PEG-Silane, MW 5000, Laysan Bio, Inc.) was used. After this incubation, the channel was thoroughly washed with T50 buffer to remove unbound BSA-biotin or biotinylated PEG. Next, a solution of NeutrAvidin (0.2 mg/mL, ThermoFisher Scientific) was flowed through the channel for 15 min; the NeutrAvidin bound to the biotinylated surface. Excess NeutrAvidin was removed by a subsequent wash with T50 buffer. Between each step, the common flow channel, also called the providing channel, was flushed with T50 buffer to avoid cross-contamination and ensure the purity of the immobilization process. All solutions were introduced at a controlled slow flow rate of 0.1 mm/min, achieved by reducing the channel back pressure to 3 psi, to maintain homogeneity and prevent material heterogeneity. Before introducing the DNA origami motors, the working channel surface was equilibrated with working buffer. For ensemble measurements, the annealed DNA origami with biotinylated anchor strands was introduced at a final concentration of 1 nM in working buffer. This solution was flowed through the channel until the desired motor density was achieved, as monitored by donor excitation laser and confirmed by the high FRET signal in the acceptor channel. Once the optimal motor density was reached, excess DNA origami-based motors were rinsed out using the working buffer. This series of steps ensured efficient and reproducible immobilization of DNA origami motors, crucial for reliable operation and data collection in downstream experiments. A comprehensive list of all immobilization commands used in the microfluidic setup is provided in **Supplementary Information Table 3**.

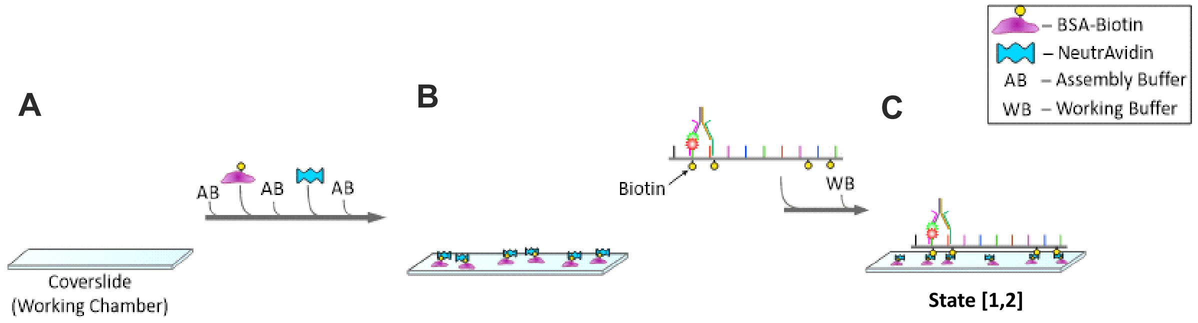

**Figure S3.** Immobilization of the motors on glass surface inside the microfluidics working chamber. **(A)** The coverslip is passivated using BSA-biotin, followed by the addition of NeutrAvidin, creating a biotin-binding surface. **(B)** The DNA origami track, carrying biotinylated staples on the side opposite the walker, is then introduced into the system. The track binds to the surface via biotin-avidin-biotin interaction, ensuring stable immobilization. **(C)** The motor immobilized in its initial state, [1,2], on the glass surface, ready for subsequent experimental operations.

### 1.9. Motor reset and initiation of walking

During origami annealing walkers may be positioned in different states (i.e., states [0,1], [1]). To ensure all motors start at state [1,2], the following procedure was used: First, AF1 was added to remove walkers in state [1] (i.e., connected only to T1, **Figure S4A**), leaving only motors that are connected to T0. Subsequently, low concentrations of F1 (100 nM) were introduced to ensure that motors that were standing on T0 stand on T0 and T1. This was followed by introduction of F2, washing and introduction of AF0 to ensure that all motors stand on T1 and T2 (i.e., state [1,2]). **Figure S4B** shows the changes in donor and acceptor emissions and FRET values that occur during these initiation steps.

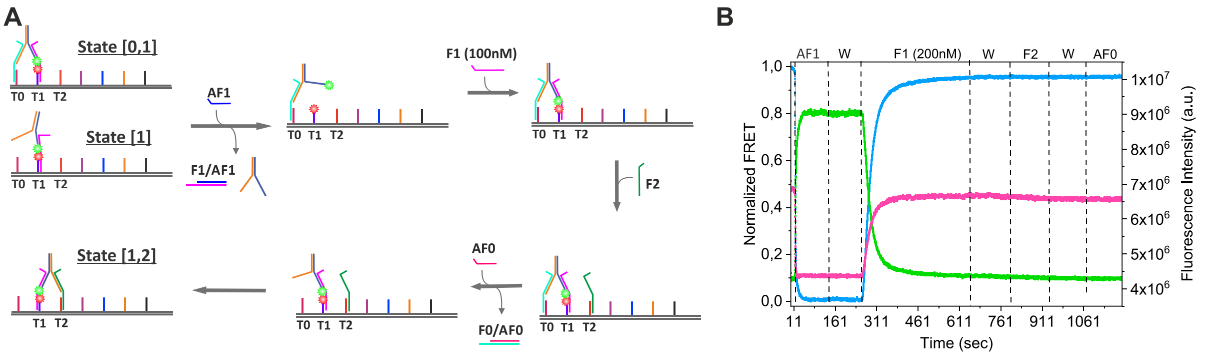

**Figure S4.** Motor initialization. **(A)** Schematic of initiation procedure. **(B)** A graph showing the FRET signal (blue) and the donor (green) and acceptor (magenta) fluorescence signals during initiation.

### 1.10. Sequence-encoded photophysics

The donor and acceptor fluorophore intensities are expected to change during motor operation because of changes in donor-acceptor distances and the FRET process. However, we noticed additional changes in donor and acceptor intensities that we attribute to two photophysical effects. The donor intensity dropped when donor-labeled L1 interacted with F5 (**Fig. S5, left panel and upper right panel**). We hypothesize that the proximity of the donor fluorophore on L1 to a guanine base in F5 led to a reduction of the donor intensity. Guanine is a well-known static quencher:[10] Its low oxidation potential facilitates photo-induced electron transfer to the excited fluorophore, producing a long-lived, dark charge-transfer state, reducing fluorescence intensity. A similar but opposite effect happened to the acceptor fluorophore when F1 was introduced. In the duplex with F1, the acceptor-labeled single-stranded T1 was no longer flexible, and this likely reduced interactions of the acceptor with the F1 guanine bases, resulting in increased acceptor intensity (**Fig. S5, bottom right panel**).

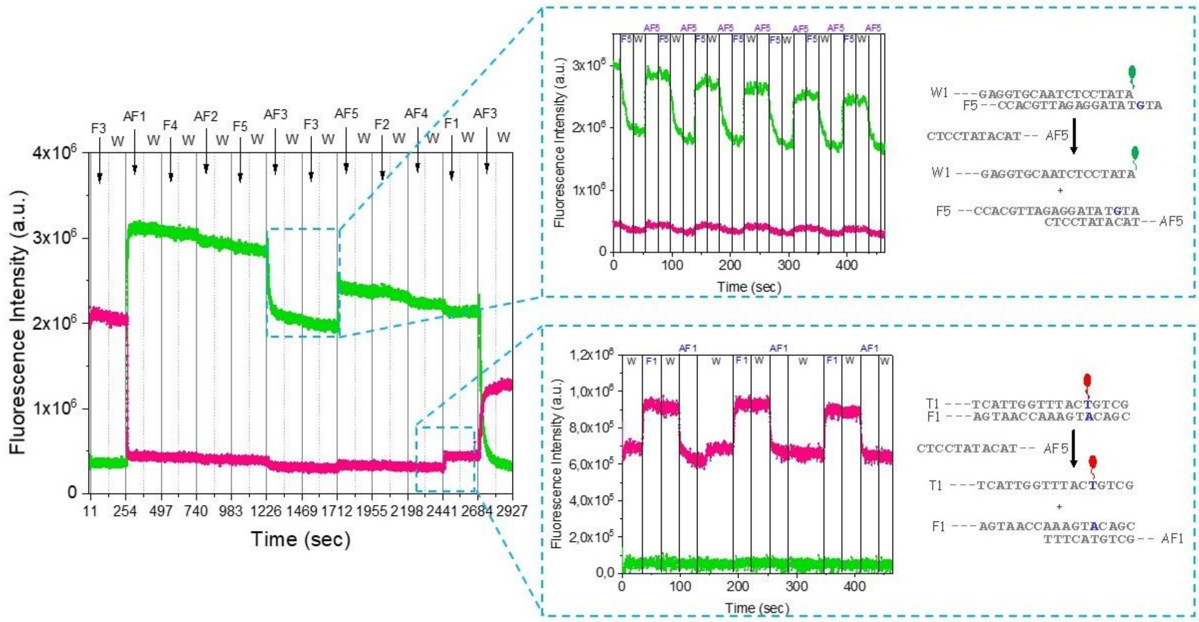

**Figure S5.** Sequence-specific alterations in dye intensities are not related to motor operation. The donor and acceptor intensities are influenced by quenching by guanine bases. Interaction of the donor-labeled L1 with F5 results in decrease donor intensity (upper right panel), and interaction of the acceptor labeled T1 with F1 results in increased acceptor intensity (lower right panel).

### 1.11. Calculation of motor operational yield

The operational yield of the bipedal motor was determined by analyzing relative changes in donor fluorescence intensities during walking (**Figure S6**). The walker leg L1 was labeled with the donor fluorophore. Donor fluorescence reflects the presence of the walker in the system. As the donor intensity is influenced by the proximity of the donor to the acceptor on foothold T1, it is therefore a reflection of motor state. To calculate the relative amount of the surviving walkers, we compared the changes in donor intensity at the beginning and at the end of each walking cycle. The walker survival rate (SR) in each cycle was calculated by dividing the change in donor fluorescence at the end of the walking cycle (upon L1 placing on T1) to the change in donor intensity in the beginning of that cycle (upon L1 lifting from T1) (**eq. 1**). This calculation provides the fraction of walkers that survived the walking cycle.

$$Eq.1: \qquad SR = \frac{GH2 - GL2}{GH1 - GL1}$$

The survival rate per step, where the yield per step is denoted Y, is calculated as shown in **eq. 2** where c is the number of steps in a cycle.

$$Eq.2: \qquad Y = SR^{\left(\frac{1}{c}\right)}$$

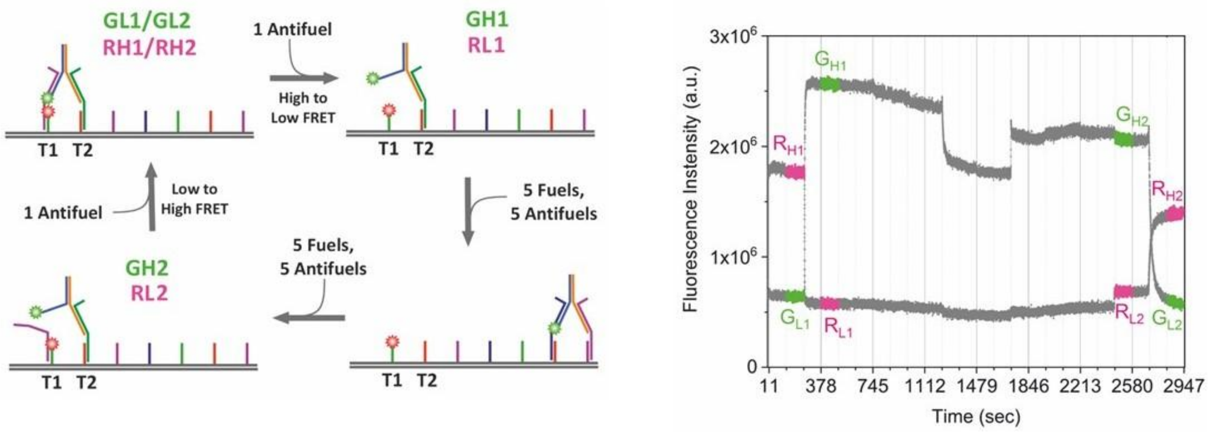

**Figure S6.** Measurement of operational yield. A schematic representation of the bipedal walker illustrates the process of lifting L1 from foothold T1 and placing of L1 to end of walking cycle. Donor fluorescence intensity over the cycle of leg lifting (GL1→GH1) to leg placing (GH2→GL2).

### 2. DNA strand sequences

**Supplementary Table 1.** Sequences of the DNA strands and labeling positions. Start and end positions refer to locations in Supplementary Table 2.

| Legs | Sequence 5'→3' |
|---|---|
| L1 | ATCAACGGCTGGCTGGTTTCTGCTCTCTAGTTTTTGAGGTGCAATCTCCTATC-ATTO550 |
| L2 | CGATGGTGTCTAGATCACTTTTTCTAGAGAGCAGAAACCAGCCAGCCGTTGAT |
| **Fuels & Antifuels** | **Sequence 5'→3'** |
| F0 (FS) | GATCTAGACACCATCGAGAGCTCGACGGTCATAGTTAGCGTTAATAGG |
| AF0 (AFS) | CCTATTAACGCTAACTATGACCGTCGAGCTCTCGATGGTG |
| F1 | CAATTTACCCAGTAACCAAAGTACAGCACTGCGATAGGAGATTGCACC |
| AF1 | CTCCTATCGCAGTGCTGTACTTTGGTTACTGGGTAAATTG |
| F2 | GATCTAGACACCATCGTTGAAACGCTACTGCCCATACAGACTGTTACG |
| AF2 | CGTAACAGTCTGTATGGGCAGTAGCGTTTCAACGATGGTG |
| F3 | ATCATTGTCGACAGAGTCCGATGTGGAAGTCAGATAGGAGATTGCACC |
| AF3 | CTCCTATCTGACTTCCACATCGGACTCTGTCGACAATGAT |
| F4 | GATCTAGACACCATCGCTCAGACCTGACGACAATGCTGGGATCAGTTA |
| AF4 | TAACTGATCCCAGCATTGTCGTCAGGTCTGAGCGATGGTG |
| F5 | TTGCGTGTATCAAGTTGCGCATGAGCTCCAGTGATAGGAGATTGCACC |
| AF5 | CTCCTATCACTGGAGCTCATGCGCAACTTGATACACGCAA |
| F6 | GATCTAGACACCATCGCTGCTAGTAATGTGCCGACTCTAGTCTGTCAA |
| AF6 | TTGACAGACTAGAGTCGGCACATTACTAGCAGCGATGGTG |

| Single Track | Start point | End point | Sequence 5'→3' |
|---|---|---|---|
| T0 (TS) | 10[79] | 8[80] | GGTGAATTTGAAATAGCAATAGCTTCAGAGGGTTTCTAACTATGACCGTCGA |
| T1 | 5[96] | 7[95] | GCTGT-ATTO647N-ACTTTGGTTACTTTTGCCAACGCGGTATTAAACCAAGTACCGGTATT |
| T2 | 3[112] | 0[112] | TGTAAATCGTTAATTTCATCTTCTAATTGAGATTTTGTATGGGCAGTAGCGT |
| T3 | 5[160] | 7[159] | TCCACATCGGACTCTGTTTTTAGGCAGATAATTTACGAGCATGTTTAAATCA |
| T4 | 3[192] | 5[191] | TTAAGACGAATGCTGATGCAAATCGTAATAAGTTTCAGCATTGTCGTCAGGT |
| T5 | 5[224] | 7[223] | AGCTCATGCGCAACTTGTTTTTCTGTCCTGCAGAACGCGCCTGTTAACGAGC |
| T6 | 6[239] | 5[255] | TCAGCTAAAGACGACGACAATAAATTGGGTAATTTAGAGTCGGCACATTACT |

| | | | |
|---|---|---|---|
| **T7** | 5[288] | 7[287] | **GCTGT-ATTO647N-ACTTTGGTTACTTTTACGACGGCCTATTACGCCAGCTGGATTGTATA** |
| **T8** | 3[320] | 5[319] | **ACGTGGACAGCAAGCGGTCCACGCATGCCTGCTTTTGTATGGGCAGTAGCGT** |
| **T9** | 5[352] | 7[351] | **TCCACATCGGACTCTGTTTTCTCGAATTAAGCGCCATTCGCCATTGTTAAAT** |
| T10 | 3[384] | 5[383] | **CATCACCCCTTTTCACCAGTGAGAAGCTGTTTTTTCAGCATTGTCGTCAGGT** |
| T11 | 5[416] | 7[415] | **AGCTCATGCGCAACTTGTTTACACAACAGCCTCAGGAAGATCGCCCTGTAGC** |
| **T12** | 3[448] | 5[447] | **GGAGCCCCATTAATGAATCGGCCATAAAGTGTTTTAGAGTCGGCACATTACT** |
| **T13** | 8[463] | 6[464] | **GCTGTACTTTGGT-ATTO647N-TACTTTTTAAAGATTTCGGATTCTCCGTGGCATCGTAA** |

| **Biotinylated strands** | **Start point** | **End point** | **Sequence 5'→3'** |
|---|---|---|---|
| Biotin-1 | 3[112] | 0[112] | **5' Biotin-TTTTTGAATAACAAATCGCGCAGAGGCGAAACCACCAGAAGGAGATTAGAG** |
| Biotin-2 | 3[432] | 0[432] | **5' Biotin-TTTTAATCGGAACGCTGCGCGTAACCACGAAATACCTACATTTGAATACGT** |
| Biotin-3 | 10[287] | 13[287] | **5' Biotin-TTTTGAGTAGATTGTTTAGACTGGATAGTTTACCAGACGACGAGGACAGAT** |
| Biotin-4 | 12[127] | 15[127] | **5' Biotin-TTTTGACAGGAGTCAGTGCCTTGAGTAACCGCCACCCTCAGAAGTCTTTCC** |
| Biotin-5 | 12[447] | 15[447] | **5' Biotin-TTTTACGTTAATGTAAATTGGGCTTGAGAAATACGTAATGCCACCCTCAGC** |

**Supplementary Table 2.** DNA origami staple strand sequences.

| **Track** | **Start point** | **End point** | **Sequence 5'→3'** |
|---|---|---|---|
| LCO-2 | 0[79] | 1[63] | AAGTATTAGACTTTACAAACAATTTTATTAAT |
| LCO-3 | 0[111] | 1[95] | CCGTCAATAGATAATACATTTGAATTTTGCG |
| LCO-4 | 0[143] | 1[127] | CTTTAGGAGCACTAACAACTAATAGCGGAATT |
| LCO-5 | 0[175] | 1[159] | GAAAGGAATTGAGGAAGGTTATCTGATGGCA |
| LCO-6 | 0[207] | 1[191] | TCAATATCTGGTCAGTTGGCAAATTTTGGATT |
| LCO-7 | 0[239] | 1[223] | CCTTGCTGAACCTCAAATATCAAAGAACCTA |
| LCO-8 | 0[271] | 0[240] | AGCCAGCAGCAAATGAAAAATCTAAAGCATCA |
| LCO-9 | 0[303] | 1[287] | TTAACACCGCCTGCAACAGTGCCACGCAAAT |
| LCO-10 | 0[335] | 1[319] | CAGAAGATAAAACAGAGGTGAGGCTGATTAGT |
| LCO-11 | 0[367] | 1[351] | TCGCCATTAAAAATACCGAACGAAGAACTCA |
| LCO-12 | 0[399] | 1[383] | TCTTTAATGCGCGAACTGATAGCCTCCAGAAC |
| LCO-13 | 0[431] | 1[415] | GGCACAGACAATATTTTTGAATGAGGAAAAA |

| | | | |
|---|---|---|---|
| LCO-14 | 0[463] | 1[447] | AACCCTTCTGACCTGAAAGCGTAATGACGCTC |
| LCO-15 | 0[487] | 1[479] | GGACATTCTGGCCAACATTGGCAG |
| LCO-16 | 1[64] | 3[63] | TTTAAAAGGATGATGAAACAAACTTTCATTT |
| LCO-17 | 1[96] | 3[95] | GAACAAAGAATTATTCATTTCAATAGTACATA |
| LCO-18 | 1[128] | 3[127] | ATCATCATGAATACCAAGTTACACTTGCTTC |
| LCO-19 | 1[160] | 3[159] | ATTCATCAGAGAAACAATAACGGATTTTCCCT |
| LCO-20 | 1[192] | 3[191] | ATACTTCTGAATATACAGTAACAAGCTTAGA |
| LCO-21 | 1[224] | 3[223] | CCATATCAAATTGCGTAGATTTTCAGTGAATT |
| LCO-22 | 1[256] | 2[240] | ACCGAGTAGTGTTTTTATAATCAAAAACAGA |
| LCO-23 | 1[288] | 3[287] | TAACCGTTTAGACAGGAACGGTACAGTGTTGT |
| LCO-24 | 1[320] | 3[319] | AATAACATGAGCTAAACAGGAGGATTAAAGA |
| LCO-25 | 1[352] | 3[351] | AACTATCGATAACGTGCTTTCCTCCGAAAAAC |
| LCO-26 | 1[384] | 3[383] | AATATTACGCGCGTACTATGGTTACGTGAAC |
| LCO-27 | 1[416] | 3[415] | CGCTCATGCACACCCGCCGCGCTTGGGTCGAG |
| LCO-28 | 1[448] | 3[447] | AATCGTCTCAAGTGTAGCGGTCACCCTAAAG |
| LCO-29 | 1[480] | 3[479] | ATTCACCAGAAAGCGAAAGGAGCGCGGGGAAA |
| LCO-30 | 2[79] | 0[80] | CAAAAGAATTTGAGTAACATTATCGGATTTAG |
| LCO-31 | 2[143] | 0[144] | ATTGCTTTATTCCTGATTATCAGATAAAATAT |
| LCO-32 | 2[175] | 0[176] | TACATCGGATATAATCCTGATTGCAACAGTT |
| LCO-33 | 2[207] | 0[208] | CGTCAGATGAATAATGGAAGGGTTACCCTCAA |
| LCO-34 | 2[239] | 1[255] | AATAAAGAAAATTATTTGCACGTGTGAGGCC |
| LCO-35 | 2[271] | 0[272] | CCTGAGAAAAAGAGTCTGTCCATCACGCTGAG |
| LCO-36 | 2[303] | 0[304] | AGGGATTTGTAGCAATACTTCTTGGTCAGTA |
| LCO-37 | 2[335] | 0[336] | CAGAGCGGCACTTGCCTGAGTAGAACCACCAG |
| LCO-38 | 2[367] | 0[368] | GAGCACGTGCCTTGCTGGTAATACTAAAACA |
| LCO-39 | 2[399] | 0[400] | GCTACAGGCGCCAGCCATTGCAACGCTATTAG |
| LCO-40 | 2[463] | 0[464] | GGCGCTGGGAAATGGATTATTTACAGAGATAG |
| LCO-41 | 3[64] | 5[63] | GAATTACCGTTAAATAAGAATAAATTAGTATC |
| LCO-42 | 3[96] | 5[95] | AATCAATATTTGAAATACCGACCAGTATAAA |
| LCO-43 | 3[112] | 0[112] | TGAATAACAAATCGCGCAGAGGCGAAACCACCAGAAGGAGATTAGAG |
| LCO-44 | 3[128] | 5[127] | TGTAAATCGTTAATTTCATCTTCTAATTGAGA |
| LCO-45 | 3[160] | 5[159] | TAGAATCCAACGCGAGAAAACTTATGTAATT |
| LCO-46 | 3[192] | 5[191] | TTAAGACGAATGCTGATGCAAATCGTAATAAG |
| LCO-47 | 3[224] | 5[223] | TATCAAAACGGCTTAGGTTGGGTTAAAGTAA |
| LCO-48 | 3[256] | 4[240] | AAGAATAGTCGGCAAAATCCCTTACTACCTTT |
| LCO-49 | 3[288] | 5[287] | TCCAGTTTGCGAAAATCCTGTTTGTTGTAAA |

| | | | |
|---|---|---|---|
| LCO-50 | 3[320] | 5[319] | ACGTGGACAGCAAGCGGTCCACGCATGCCTGC |
| LCO-51 | 3[352] | 5[351] | CGTCTATCGCCCTTCACCGCCTGGTACCGAG |
| LCO-52 | 3[384] | 5[383] | CATCACCCCTTTTCACCAGTGAGAAGCTGTTT |
| LCO-53 | 3[416] | 5[415] | GTGCCGTAGGTTTGCGTATTGGGACAATTCC |
| LCO-54 | 3[432] | 0[432] | AATCGGAACGCTGCGCGTAACCACGAAATACCTACATTTGAATACGT |
| LCO-55 | 3[448] | 5[447] | GGAGCCCCATTAATGAATCGGCCATAAAGTGT |
| LCO-56 | 3[480] | 5[479] | GCCGGCGATTTCCAGTCGGGAAACAGCTAACT |
| LCO-57 | 4[79] | 2[80] | AATAAGGCTTTTTAATGGAAACTACCTGAG |
| LCO-58 | 4[143] | 2[144] | ATATTTTAGTCGCTATTAATTAATTCGCCTG |
| LCO-59 | 4[175] | 2[176] | AGACAAAGTTGAAAACATAGCGATGTACCTTT |
| LCO-60 | 4[207] | 2[208] | TATATGTACTGAGAAGAGTCAATAGGTTTAA |
| LCO-61 | 4[239] | 3[255] | TTAACCTCTCATAGGTCTGAGAGATAAATCAA |
| LCO-62 | 4[271] | 2[272] | TTCCGAAACCCGAGATAGGGTTGGCCAGAAT |
| LCO-63 | 4[303] | 2[304] | CCCAGCAGGGAACAAGAGTCCACTCCGATTAA |
| LCO-64 | 4[335] | 2[336] | AGAGTTGCTCCAACGTCAAAGGGGTTAGAAT |
| LCO-65 | 4[367] | 2[368] | AGCTGATTAGGGCGATGGCCCACTGCTTTGAC |
| LCO-66 | 4[399] | 2[400] | TGGTTTTTAAATCAAGTTTTTTGAATGCGCC |
| LCO-67 | 4[463] | 2[464] | CCAGCTGCCGATTTAGAGCTTGAGGCGCTAG |
| LCO-68 | 5[64] | 7[63] | ATATGCGTCAAGCAAGCCGTTTTATAGCAAG |
| LCO-69 | 5[96] | 7[95] | GCCAACGCGGTATTAAACCAAGTACCGGTATT |
| LCO-70 | 5[128] | 7[127] | ATCGCCATTCGGCTGTCTTTCCTAACCTCCC |
| LCO-71 | 5[160] | 7[159] | TAGGCAGATAATTTACGAGCATGTTTAAATCA |
| LCO-72 | 5[192] | 7[191] | AGAATATAAGTCCTGAACAAGAAGCTACAAT |
| LCO-73 | 5[224] | 7[223] | TTCTGTCCTGCAGAACGCGCCTGTTAACGAGC |
| LCO-74 | 5[256] | 6[240] | CGCCAGGGTGCAAGGCGATTAAGCAACATGT |
| LCO-75 | 5[288] | 7[287] | ACGACGGCCTATTACGCCAGCTGGATTGTATA |
| LCO-76 | 5[320] | 7[319] | AGGTCGACTTGGGAAGGGCGATCTAATATTT |
| LCO-77 | 5[352] | 7[351] | CTCGAATTAAGCGCCATTCGCCATTGTTAAAT |
| LCO-78 | 5[384] | 7[383] | CCTGTGTGCGGCACCGCTTCTGGACGCCATC |
| LCO-79 | 5[416] | 7[415] | ACACAACAGCCTCAGGAAGATCGCCCTGTAGC |
| LCO-80 | 5[448] | 7[447] | AAAGCCTGCTGCCAGTTTGAGGGAGCGAGTA |
| LCO-81 | 5[480] | 7[479] | CACATTAATTGGTGTAGATGGGCGGAACAAAC |
| LCO-82 | 6[79] | 4[80] | ATCGAGAATATACAAATTCTTACCGTGTGATA |
| LCO-83 | 6[111] | 3[111] | CAAGAACGTCAACAGTAGGGCTTGACCTAAATTTAATGGTATGTGAG |
| LCO-84 | 6[143] | 4[144] | ATCAATAAATTTAACAACGCCAACTTTCAAAT |
| LCO-85 | 6[175] | 4[176] | TCCCATCCGGCATTTTCGAGCCACAATCGCA |

| | | | |
|---|---|---|---|
| LCO-86 | 6[207] | 4[208] | AATAGATAAAGTACCGACAAAAGGTATATAAC |
| LCO-87 | 6[239] | 5[255] | TCAGCTAAAGACGACGACAATAAATTGGGTAA |
| LCO-88 | 6[271] | 4[272] | GGATGTGCTTTTCCCAGTCACGACGATGGTGG |
| LCO-89 | 6[303] | 4[304] | CCTCTTCGCAGTGCCAAGCTTGCTGGTTTGC |
| LCO-90 | 6[335] | 4[336] | CGCAACTGTCTAGAGGATCCCCGGGCCCTGAG |
| LCO-91 | 6[367] | 4[368] | ACCAGGCACGTAATCATGGTCATCGGGCAAC |
| LCO-92 | 6[399] | 4[400] | CAGCTTTCAAATTGTTATCCGCTCCGCCAGGG |
| LCO-93 | 6[431] | 3[431] | CAGTATCGTACGAGCCGGAAGCAACGCGCGGGGAGAGGCAAGCACTA |
| LCO-94 | 6[463] | 4[464] | CCGTGCATGGGTGCCTAATGAGTGCTGTCGTG |
| LCO-95 | 7[64] | 9[63] | CAAATCAGCGCTAATATCAGAGAGTAAGAGCA |
| LCO-96 | 7[96] | 9[95] | CTAAGAACACACCCTGAACAAAGATCTTACC |
| LCO-97 | 7[128] | 9[127] | GACTTGCGAGGGAAGCGCATTAGAGCAGATAG |
| LCO-98 | 7[160] | 9[159] | AGATTAGTGCAGCCTTTACAGAGCCGAGGAA |
| LCO-99 | 7[192] | 9[191] | TTTATCCTCGATTTTTTGTTTAACAAAGAACT |
| LCO-100 | 7[224] | 9[223] | GTCTTTCCCCATATTATTTATCCGCAGTATG |
| LCO-101 | 7[256] | 8[240] | AGAAAAGCTCATATGTACCCCGGTGTTACAAA |
| LCO-102 | 7[288] | 10[288] | AGCAAATAATGAACGGTAATCGTGGGGCGCGAGCTGAAACTGCGAAC |
| LCO-103 | 7[320] | 9[319] | TGTTAAAATGCCTGAGAGTCTGGATACTAATA |
| LCO-104 | 7[352] | 9[351] | CAGCTCATTTTGAGAGATCTACATCATACAG |
| LCO-105 | 7[384] | 9[383] | AAAAATAACTGATAAATTAATGCCTAAGCAAT |
| LCO-106 | 7[416] | 9[415] | CAGCTTTCTCACCATCAATATGATCGGTTGT |
| LCO-107 | 7[448] | 9[447] | ACAACCCGCAAAAGGGTGAGAAAGATACTTTT |
| LCO-108 | 7[480] | 9[479] | GGCGGATTTGCAATGCCTGAGTAACAAGGATA |
| LCO-109 | 8[79] | 6[80] | TAATTGAGATATAGAAGGCTTATCCGCACTC |
| LCO-110 | 8[111] | 6[112] | TTAACTGAGCGAGGCGTTTTAGCGTATCATTC |
| LCO-111 | 8[143] | 6[144] | ATAAAAACGGAGGTTTTGAAGCCAGAAACCA |
| LCO-112 | 8[175] | 6[176] | TGAAAATATGCTATTTTGCACCCAAAATAATA |
| LCO-113 | 8[207] | 6[208] | ATAAGAAAGAATCTTACCAACGCTTATCAAC |
| LCO-114 | 8[239] | 7[255] | ATAAACAGAGAGCCTAATTTGCCATGATAATC |
| LCO-115 | 8[271] | 6[272] | CATGTCAACCCAAAAACAGGAAGCGAAAGGG |
| LCO-116 | 8[303] | 6[304] | GAGAATCGTTTAAATTGTAAACGTGGTGCGGG |
| LCO-117 | 8[335] | 6[336] | CAGGTCATTTCGCATTAAATTTTTCAGGCTG |
| LCO-118 | 8[367] | 6[368] | TAGCTATTTTTTTAACCAATAGGATGCCGGAA |
| LCO-119 | 8[399] | 6[400] | CGTTCTAGTTCGCGTCTGGCCTTACTCCAGC |
| LCO-120 | 8[431] | 6[432] | CAGTCAAAATCAACATTAAATGTGGACGACGA |

| | | | |
|---|---|---|---|
| LCO-121 | 8[463] | 6[464] | TAAAGATTTCGGATTCTCCGTGGCATCGTAA |
| LCO-122 | 9[64] | 11[63] | AGAAACAAATCACCGTCACCGACCACCAGTA |
| LCO-123 | 9[96] | 11[95] | GAAGCCCTTATTGACGGAAATTATGGAAACGT |
| LCO-124 | 9[128] | 12[128] | CCGAACAAACATTCAACCGATTGCACCGTAATCAGTAGCCCAGCATT |
| LCO-125 | 9[160] | 11[159] | ACGCAATATATGGTTTACCAGCGCGCCTTTAG |
| LCO-126 | 9[192] | 11[191] | GGCATGATGTTTATTTTGTCACACGGCATTT |
| LCO-127 | 9[224] | 11[223] | TTAGCAAATATAAAAGAAACGCAAGTTTGCCA |
| LCO-128 | 9[256] | 10[240] | ACCTGTTTATACATTTCGCAAATCATAAAGG |
| LCO-129 | 9[320] | 11[319] | GTAGTAGCTTTCATTCCATATAAATCGTCAT |
| LCO-130 | 9[352] | 11[351] | GCAAGGCATAAATATGCAACTAAAAATGCTTT |
| LCO-131 | 9[384] | 11[383] | AAAGCCTCGCTGAATATAATGCTCCATAAAT |
| LCO-132 | 9[416] | 11[415] | ACCAAAAAGGTCATTTTTGCGGATCTATTATA |
| LCO-133 | 9[448] | 12[448] | GCGGGAGAAGAGTACCTTTAATTAAAAAGATTAAGAGGAAAAAATCT |
| LCO-134 | 9[480] | 11[479] | AAAATTTTACCGGAAGCAAACTCCAAATATCG |
| LCO-135 | 10[79] | 8[80] | GGTGAATTTGAAATAGCAATAGCTTCAGAGGG |
| LCO-136 | 10[111] | 8[112] | AAGGTAAATTTTAAGAAAAGTAACGGGAGAA |
| LCO-137 | 10[143] | 8[144] | AAAGGGCGAGTTACCAGAAGGAAAAGAATAAC |
| LCO-138 | 10[175] | 8[176] | AAAATTCAATAACGGAATACCCAGTCAAAAA |
| LCO-139 | 10[207] | 8[208] | CGGAATAATAAGACTCCTTATTACCAATCCAA |
| LCO-140 | 10[239] | 9[255] | TGGCAACACGTAGAAAATACATAGGTCAATA |
| LCO-141 | 10[271] | 8[272] | ACCATTAGAGCTATATTTTCATTTAAAACTAG |
| LCO-142 | 10[287] | 13[287] | GAGTAGATTGTTTAGACTGGATAGTTTACCAGACGACGAGGACAGAT |
| LCO-143 | 10[303] | 8[304] | TCCCAATTAGGTGGCATCAATTCGCAAACAA |
| LCO-144 | 10[335] | 8[336] | TCTGGAAGATTAACATCCAATAAAAAGGCTAT |
| LCO-145 | 10[367] | 8[368] | ACATGTTTAAGAATTAGCAAAATGGAGAGGG |
| LCO-146 | 10[399] | 8[400] | GCTTAATTAGAGCATAAAGCTAAATATTCAAC |
| LCO-147 | 10[431] | 8[432] | TGATAAGACATTATGACCCTGTAGCCGGAGA |
| LCO-148 | 10[463] | 8[464] | AGGATTAGAGCCTTTATTTCAACGTGTGTAGG |
| LCO-149 | 11[64] | 13[63] | GCACCATTTCCTCATTAAAGCCAGCAGTAAGC |
| LCO-150 | 11[96] | 13[95] | CACCAATGCGATTGGCCTTGATAAGGAGTGT |
| LCO-151 | 11[160] | 13[159] | CGTCAGACGCCGCCACCAGAACCAGTTAATG |
| LCO-152 | 11[192] | 13[191] | TCGGTCATGAACCGCCACCCTCAGTTATTCTG |
| LCO-153 | 11[224] | 13[223] | TCTTTTCAGGAACCGCCTCCCTCAGACTCCT |
| LCO-154 | 11[256] | 12[240] | TGCCAGAGGAGAGGCTTTTGCAAAAACCAGAG |
| LCO-155 | 11[320] | 13[319] | AAATATTCGCATAGTAAGAGCAACTAGGCTGG |
| LCO-156 | 11[352] | 13[351] | AAACAGTTGCAGATACATAACGCGACAAGAA |

| | | | |
|---|---|---|---|
| LCO-157 | 11[384] | 13[383] | CAAAAATCGTTGAGATTTAGGAATACGTAACA |
| LCO-158 | 11[416] | 13[415] | GTCAGAAGACAACATTATTACAGCTTGCCCT |
| LCO-159 | 11[480] | 13[479] | CGTTTTAATGGCTCATTATACCAGTTTAATCA |
| LCO-160 | 12[79] | 10[80] | CAAATAAAACCATTAGCAAGGCCTCATTAAA |
| LCO-161 | 12[111] | 10[112] | AGGTCAGAAAACCATCGATAGCAGAGGGAGGG |
| LCO-162 | 12[127] | 15[127] | GACAGGAGTCAGTGCCTTGAGTAACCGCCACCCTCAGAAGTCTTTCC |
| LCO-163 | 12[143] | 10[144] | AGCCGCCGGACAGAATCAAGTTTCAAAGACA |
| LCO-164 | 12[175] | 10[176] | CCCTCAGATGTAGCGCGTTTTCATATCAATAG |
| LCO-165 | 12[207] | 10[208] | CACCCTCAAGCCCCCTTATTAGCAGACACCA |
| LCO-166 | 12[239] | 11[255] | CCACCACCTAATCAAAATCACCGGAGAAGTTT |
| LCO-167 | 12[271] | 10[272] | AAAATAGCGGGGTAATAGTAAAATTAGTTTG |
| LCO-168 | 12[303] | 10[304] | AACCCTCGCGTCCAATACTGCGGACAGTTGAT |
| LCO-169 | 12[335] | 10[336] | ATTACGAGATTGAATCCCCCTCAGTACGGTG |
| LCO-170 | 12[367] | 10[368] | CAACTAATCAGAAAACGAGAATGAGTAGCTCA |
| LCO-171 | 12[399] | 10[400] | ATTCATCAAGGTCTTTACCCTGAGGCTTAGA |
| LCO-172 | 12[431] | 10[432] | CTAACGGACAAAGCGGATTGCATCGCTCCTTT |
| LCO-173 | 12[447] | 15[447] | ACGTTAATGTAAATTGGGCTTGAGAAATACGTAATGCCACCCTCAGC |
| LCO-174 | 12[463] | 10[464] | TTGGGAAGAGCCCGAAAGACTTCAACAGGTC |
| LCO-175 | 13[64] | 15[63] | GTCATACAATAGGAACCCATGTATACAACGC |
| LCO-176 | 13[96] | 15[95] | ACTGGTAACACCACCCTCATTTTCTCATAGTT |
| LCO-177 | 13[160] | 15[159] | CCCCCTGCACTCAGGAGGTTTAGTGTATGGGA |
| LCO-178 | 13[192] | 15[191] | AAACATGATAAGTATAGCCCGGATTCAGCGG |
| LCO-179 | 13[224] | 15[223] | CAAGAGAACAGGCGGATAAGTGCCAAAGGAAT |
| LCO-180 | 13[256] | 14[240] | ACCGAACTCGCAGACGGTCAATCGGGTTTTG |
| LCO-181 | 13[288] | 15[287] | GAACGGTGGACCTGCTCCATGTTATTGTATCG |
| LCO-182 | 13[320] | 15[319] | CTGACCTTTCATCGCCTGATAAAAATTTCTT |
| LCO-183 | 13[352] | 15[351] | CCGGATATAAGCGCGAAACAAAGTCGCCGACA |
| LCO-184 | 13[384] | 15[383] | AAGCTGCTACACTCATCTTTGACTAACCGAT |
| LCO-185 | 13[416] | 15[415] | GACGAGAACTAAAACGAAAGAGGCGGGAGTTA |
| LCO-186 | 13[480] | 15[487] | TTGTGAATTTTTTCATGAGGAAGTGGGTAGCAACGGCTAC |
| LCO-187 | 14[79] | 12[80] | CAAGCCCATGGCTTTTGATGATACTTCACAAA |
| LCO-188 | 14[111] | 12[112] | CTCAGAGCTAAGTTTTAACGGGGGTTGAGGC |
| LCO-189 | 14[143] | 12[144] | CCTCAGAACAGTGCCCGTATAAACACCACCAG |
| LCO-190 | 14[175] | 12[176] | ATCACCGTCTATTTCGGAACCTAAGCCACCA |
| LCO-191 | 14[207] | 12[208] | GGTTGATAAAGTATTAAGAGGCTGAGAGCCGC |
| LCO-192 | 14[239] | 13[255] | CTCAGTACGGATTAGGATTAGCGATAAGGGA |

| | | | |
|---|---|---|---|
| LCO-193 | 14[271] | 12[272] | GAACGAGGGACCAACTTTGAAAGATAAAAACC |
| LCO-194 | 14[303] | 12[304] | AAATCCGCTACAGACCAGGCGCAACTATCAT |
| LCO-195 | 14[335] | 12[336] | GATTTGTACATCAAGAGTAATCTTCAAAAGGA |
| LCO-196 | 14[367] | 12[368] | ATTATACCTCATTACCCAAATCAACCACATT |
| LCO-197 | 14[399] | 12[400] | ACACTAAACATTCAGTGAATAAGGGTAGAAAG |
| LCO-198 | 14[431] | 12[432] | GCACCAACACACCAGAACGAGTAAAAACGAA |
| LCO-199 | 14[463] | 12[464] | AACGGGTAATGGTTTAATTTCAACTCAGGACG |
| LCO-200 | 15[64] | 14[80] | CTGTAGCATTCCACAGACAGCCCAGGGATAG |
| LCO-201 | 15[96] | 14[112] | AGCGTAACGATCTAAAGTTTTGTCCCGCCACC |
| LCO-202 | 15[128] | 14[144] | AGACGTTAGTAAATGAATTTTCTACCGCCAC |
| LCO-203 | 15[160] | 14[176] | TTTTGCTAAACAACTTTCAACAGTATAGGTGT |
| LCO-204 | 15[192] | 14[208] | AGTGAGAATAGAAAGGAACAACTGTCGAGAG |
| LCO-205 | 15[224] | 15[255] | TGCGAATAATAATTTTGAAAATCTCCAAAAAA |
| LCO-206 | 15[256] | 14[272] | AAGGCTCCAAAAGGAGCCTTTAACTTAGCCG |
| LCO-207 | 15[288] | 14[304] | GTTTATCAGCTTGCTTTCGAGGTGTTGTGTCG |
| LCO-208 | 15[320] | 14[336] | AAACAGCTTGATACCGATAGTTGACAACGGA |
| LCO-209 | 15[352] | 14[368] | ATGACAACAACCATCGCCCACGCACCCCAGCG |
| LCO-210 | 15[384] | 14[400] | ATATTCGGTCGCTGAGGCTTGCAAAAAGAAT |
| LCO-211 | 15[416] | 14[432] | AAGGCCGCTTTTGCGGGATCGTCACTACGAAG |
| LCO-212 | 15[448] | 14[464] | AGCGAAAGACAGCATCGGAACGATTCCATTA |

### 3. Computer control of microfluidics

**Supplementary Table 3.** Microfluidics commands and time durations for motor immobilization commands.

| Command No. | Command | Time (seconds) | Washing Buffer | AF1 | F1 | AF2 | F2 | AF3 | F3 | AF4 | F4 | AF5 | F5 | AF6 | F6 | | | | CH1 | CH2 | CH3 | CH4 | CH5 | CH6 | WASTE | EXIT | Flooring Buffer | BSA-Biotin | NeutrAvidin |
|---|---|---|---|---|---|---|---|---|---|---|---|---|---|---|---|---|---|---|---|---|---|---|---|---|---|---|---|---|---|
| 1 | wash waste | 100 | 0 | 0 | 0 | 0 | 0 | 0 | 0 | 0 | 0 | 0 | 0 | 0 | 0 | 0 | 0 | 0 | 0 | 0 | 0 | 0 | 0 | 0 | 1 | 0 | 1 | 0 | 0 |
| 2 | wash | 300 | 0 | 0 | 0 | 0 | 0 | 0 | 0 | 0 | 0 | 0 | 0 | 0 | 0 | 0 | 0 | 0 | 0 | 0 | 0 | 0 | 0 | 1 | 0 | 1 | 1 | 0 | 0 |
| 3 | BSA to waste | 100 | 0 | 0 | 0 | 0 | 0 | 0 | 0 | 0 | 0 | 0 | 0 | 0 | 0 | 0 | 0 | 0 | 0 | 0 | 0 | 0 | 0 | 0 | 1 | 0 | 0 | 1 | 0 |
| 4 | BSA | 900 | 0 | 0 | 0 | 0 | 0 | 0 | 0 | 0 | 0 | 0 | 0 | 0 | 0 | 0 | 0 | 0 | 0 | 0 | 0 | 0 | 0 | 1 | 0 | 1 | 0 | 1 | 0 |
| 5 | wash waste | 100 | 0 | 0 | 0 | 0 | 0 | 0 | 0 | 0 | 0 | 0 | 0 | 0 | 0 | 0 | 0 | 0 | 0 | 0 | 0 | 0 | 0 | 0 | 1 | 0 | 1 | 0 | 0 |
| 6 | wash | 300 | 0 | 0 | 0 | 0 | 0 | 0 | 0 | 0 | 0 | 0 | 0 | 0 | 0 | 0 | 0 | 0 | 0 | 0 | 0 | 0 | 0 | 1 | 0 | 1 | 1 | 0 | 0 |

| | | | | | | | | | | | | | | | | | | | | | | | | | | | | | |
|---|---|---|---|---|---|---|---|---|---|---|---|---|---|---|---|---|---|---|---|---|---|---|---|---|---|---|---|---|---|
| 7 | NeutrAvidin to waste | 10<br>0 | 0 | 0 | 0 | 0 | 0 | 0 | 0 | 0 | 0 | 0 | 0 | 0 | 0 | 0 | 0 | 0 | 0 | 0 | 0 | 0 | 0 | 0 | 1 | 0 | 0 | 0 | 1 |
| 8 | NeutrAvidin | 90<br>0 | 0 | 0 | 0 | 0 | 0 | 0 | 0 | 0 | 0 | 0 | 0 | 0 | 0 | 0 | 0 | 0 | 0 | 0 | 0 | 0 | 0 | 1 | 0 | 1 | 0 | 0 | 1 |
| 9 | wash waste | 10<br>0 | 0 | 0 | 0 | 0 | 0 | 0 | 0 | 0 | 0 | 0 | 0 | 0 | 0 | 0 | 0 | 0 | 0 | 0 | 0 | 0 | 0 | 0 | 1 | 0 | 1 | 0 | 0 |
| 10 | wash | 30<br>0 | 0 | 0 | 0 | 0 | 0 | 0 | 0 | 0 | 0 | 0 | 0 | 0 | 0 | 0 | 0 | 0 | 0 | 0 | 0 | 0 | 0 | 1 | 0 | 1 | 1 | 0 | 0 |

**Supplementary Table 4.** Microfluidics commands and time durations for a walking experiment with 30 seconds wash time and 60 seconds fuel/antifuel introduction.

| Command No. | Command | Time(seconds) | Washing Buffer | AF1 | F1 | AF2 | F2 | AF3 | F3 | AF4 | F4 | AF5 | F5 | AF6 | F6 | | | | CH1 | CH2 | CH3 | CH4 | CH5 | CH6 | WASTE | EXIT | Flooring Buffer | BSA-Biotin | NeutrAvidin |
|---|---|---|---|---|---|---|---|---|---|---|---|---|---|---|---|---|---|---|---|---|---|---|---|---|---|---|---|---|---|
| 1 | wash | 30 | 1 | 0 | 0 | 0 | 0 | 0 | 0 | 0 | 0 | 0 | 0 | 0 | 0 | 0 | 0 | 0 | 0 | 1 | 0 | 0 | 0 | 0 | 0 | 1 | 0 | 0 | 0 |
| 2 | F3 | 60 | 0 | 0 | 0 | 0 | 0 | 0 | 1 | 0 | 0 | 0 | 0 | 0 | 0 | 0 | 0 | 0 | 0 | 1 | 0 | 0 | 0 | 0 | 0 | 1 | 0 | 0 | 0 |
| 3 | wash | 30 | 1 | 0 | 0 | 0 | 0 | 0 | 0 | 0 | 0 | 0 | 0 | 0 | 0 | 0 | 0 | 0 | 0 | 1 | 0 | 0 | 0 | 0 | 0 | 1 | 0 | 0 | 0 |
| 4 | AF1 | 60 | 0 | 1 | 0 | 0 | 0 | 0 | 0 | 0 | 0 | 0 | 0 | 0 | 0 | 0 | 0 | 0 | 0 | 1 | 0 | 0 | 0 | 0 | 0 | 1 | 0 | 0 | 0 |
| 5 | wash | 30 | 1 | 0 | 0 | 0 | 0 | 0 | 0 | 0 | 0 | 0 | 0 | 0 | 0 | 0 | 0 | 0 | 0 | 1 | 0 | 0 | 0 | 0 | 0 | 1 | 0 | 0 | 0 |
| 6 | F4 | 60 | 0 | 0 | 0 | 0 | 0 | 0 | 0 | 0 | 1 | 0 | 0 | 0 | 0 | 0 | 0 | 0 | 0 | 1 | 0 | 0 | 0 | 0 | 0 | 1 | 0 | 0 | 0 |
| 7 | wash | 30 | 1 | 0 | 0 | 0 | 0 | 0 | 0 | 0 | 0 | 0 | 0 | 0 | 0 | 0 | 0 | 0 | 0 | 1 | 0 | 0 | 0 | 0 | 0 | 1 | 0 | 0 | 0 |
| 8 | AF2 | 60 | 0 | 0 | 0 | 1 | 0 | 0 | 0 | 0 | 0 | 0 | 0 | 0 | 0 | 0 | 0 | 0 | 0 | 1 | 0 | 0 | 0 | 0 | 0 | 1 | 0 | 0 | 0 |
| 9 | wash | 30 | 1 | 0 | 0 | 0 | 0 | 0 | 0 | 0 | 0 | 0 | 0 | 0 | 0 | 0 | 0 | 0 | 0 | 1 | 0 | 0 | 0 | 0 | 0 | 1 | 0 | 0 | 0 |
| 10 | F5 | 60 | 0 | 0 | 0 | 0 | 0 | 0 | 0 | 0 | 0 | 0 | 1 | 0 | 0 | 0 | 0 | 0 | 0 | 1 | 0 | 0 | 0 | 0 | 0 | 1 | 0 | 0 | 0 |
| 11 | wash | 30 | 1 | 0 | 0 | 0 | 0 | 0 | 0 | 0 | 0 | 0 | 0 | 0 | 0 | 0 | 0 | 0 | 0 | 1 | 0 | 0 | 0 | 0 | 0 | 1 | 0 | 0 | 0 |
| 12 | AF3 | 60 | 0 | 0 | 0 | 0 | 0 | 1 | 0 | 0 | 0 | 0 | 0 | 0 | 0 | 0 | 0 | 0 | 0 | 1 | 0 | 0 | 0 | 0 | 0 | 1 | 0 | 0 | 0 |
| 13 | wash | 30 | 1 | 0 | 0 | 0 | 0 | 0 | 0 | 0 | 0 | 0 | 0 | 0 | 0 | 0 | 0 | 0 | 0 | 1 | 0 | 0 | 0 | 0 | 0 | 1 | 0 | 0 | 0 |
| 14 | F6 | 60 | 0 | 0 | 0 | 0 | 0 | 0 | 0 | 0 | 0 | 0 | 0 | 0 | 1 | 0 | 0 | 0 | 0 | 1 | 0 | 0 | 0 | 0 | 0 | 1 | 0 | 0 | 0 |
| 15 | wash | 30 | 1 | 0 | 0 | 0 | 0 | 0 | 0 | 0 | 0 | 0 | 0 | 0 | 0 | 0 | 0 | 0 | 0 | 1 | 0 | 0 | 0 | 0 | 0 | 1 | 0 | 0 | 0 |
| 16 | AF4 | 60 | 0 | 0 | 0 | 0 | 0 | 0 | 0 | 1 | 0 | 0 | 0 | 0 | 0 | 0 | 0 | 0 | 0 | 1 | 0 | 0 | 0 | 0 | 0 | 1 | 0 | 0 | 0 |
| 17 | wash | 30 | 1 | 0 | 0 | 0 | 0 | 0 | 0 | 0 | 0 | 0 | 0 | 0 | 0 | 0 | 0 | 0 | 0 | 1 | 0 | 0 | 0 | 0 | 0 | 1 | 0 | 0 | 0 |
| 18 | F1 | 60 | 0 | 0 | 1 | 0 | 0 | 0 | 0 | 0 | 0 | 0 | 0 | 0 | 0 | 0 | 0 | 0 | 0 | 1 | 0 | 0 | 0 | 0 | 0 | 1 | 0 | 0 | 0 |
| 19 | wash | 30 | 1 | 0 | 0 | 0 | 0 | 0 | 0 | 0 | 0 | 0 | 0 | 0 | 0 | 0 | 0 | 0 | 0 | 1 | 0 | 0 | 0 | 0 | 0 | 1 | 0 | 0 | 0 |
| 20 | AF5 | 60 | 0 | 0 | 0 | 0 | 0 | 0 | 0 | 0 | 0 | 1 | 0 | 0 | 0 | 0 | 0 | 0 | 0 | 1 | 0 | 0 | 0 | 0 | 0 | 1 | 0 | 0 | 0 |
| 21 | wash | 30 | 1 | 0 | 0 | 0 | 0 | 0 | 0 | 0 | 0 | 0 | 0 | 0 | 0 | 0 | 0 | 0 | 0 | 1 | 0 | 0 | 0 | 0 | 0 | 1 | 0 | 0 | 0 |
| 22 | F5 | 60 | 0 | 0 | 0 | 0 | 0 | 0 | 0 | 0 | 0 | 0 | 1 | 0 | 0 | 0 | 0 | 0 | 0 | 1 | 0 | 0 | 0 | 0 | 0 | 1 | 0 | 0 | 0 |
| 23 | wash | 30 | 1 | 0 | 0 | 0 | 0 | 0 | 0 | 0 | 0 | 0 | 0 | 0 | 0 | 0 | 0 | 0 | 0 | 1 | 0 | 0 | 0 | 0 | 0 | 1 | 0 | 0 | 0 |
| 24 | AF1 | 60 | 0 | 1 | 0 | 0 | 0 | 0 | 0 | 0 | 0 | 0 | 0 | 0 | 0 | 0 | 0 | 0 | 0 | 1 | 0 | 0 | 0 | 0 | 0 | 1 | 0 | 0 | 0 |
| 25 | wash | 30 | 1 | 0 | 0 | 0 | 0 | 0 | 0 | 0 | 0 | 0 | 0 | 0 | 0 | 0 | 0 | 0 | 0 | 1 | 0 | 0 | 0 | 0 | 0 | 1 | 0 | 0 | 0 |
| 26 | F4 | 60 | 0 | 0 | 0 | 0 | 0 | 0 | 0 | 0 | 1 | 0 | 0 | 0 | 0 | 0 | 0 | 0 | 0 | 1 | 0 | 0 | 0 | 0 | 0 | 1 | 0 | 0 | 0 |
| 27 | wash | 30 | 1 | 0 | 0 | 0 | 0 | 0 | 0 | 0 | 0 | 0 | 0 | 0 | 0 | 0 | 0 | 0 | 0 | 1 | 0 | 0 | 0 | 0 | 0 | 1 | 0 | 0 | 0 |
| 28 | AF6 | 60 | 0 | 0 | 0 | 0 | 0 | 0 | 0 | 0 | 0 | 0 | 0 | 1 | 0 | 0 | 0 | 0 | 0 | 1 | 0 | 0 | 0 | 0 | 0 | 1 | 0 | 0 | 0 |
| 29 | wash | 30 | 1 | 0 | 0 | 0 | 0 | 0 | 0 | 0 | 0 | 0 | 0 | 0 | 0 | 0 | 0 | 0 | 0 | 1 | 0 | 0 | 0 | 0 | 0 | 1 | 0 | 0 | 0 |
| 30 | F3 | 60 | 0 | 0 | 0 | 0 | 0 | 0 | 1 | 0 | 0 | 0 | 0 | 0 | 0 | 0 | 0 | 0 | 0 | 1 | 0 | 0 | 0 | 0 | 0 | 1 | 0 | 0 | 0 |
| 31 | wash | 30 | 1 | 0 | 0 | 0 | 0 | 0 | 0 | 0 | 0 | 0 | 0 | 0 | 0 | 0 | 0 | 0 | 0 | 1 | 0 | 0 | 0 | 0 | 0 | 1 | 0 | 0 | 0 |
| 32 | AF5 | 60 | 0 | 0 | 0 | 0 | 0 | 0 | 0 | 0 | 0 | 1 | 0 | 0 | 0 | 0 | 0 | 0 | 0 | 1 | 0 | 0 | 0 | 0 | 0 | 1 | 0 | 0 | 0 |

| | | | | | | | | | | | | | | | | | | | | | | | | | | | | | |
|---|---|---|---|---|---|---|---|---|---|---|---|---|---|---|---|---|---|---|---|---|---|---|---|---|---|---|---|---|---|
| 33 | wash | 30 | 1 | 0 | 0 | 0 | 0 | 0 | 0 | 0 | 0 | 0 | 0 | 0 | 0 | 0 | 0 | 0 | 0 | 1 | 0 | 0 | 0 | 0 | 0 | 1 | 0 | 0 | 0 |
| 34 | F2 | 60 | 0 | 0 | 0 | 0 | 1 | 0 | 0 | 0 | 0 | 0 | 0 | 0 | 0 | 0 | 0 | 0 | 0 | 1 | 0 | 0 | 0 | 0 | 0 | 1 | 0 | 0 | 0 |
| 35 | wash | 30 | 1 | 0 | 0 | 0 | 0 | 0 | 0 | 0 | 0 | 0 | 0 | 0 | 0 | 0 | 0 | 0 | 0 | 1 | 0 | 0 | 0 | 0 | 0 | 1 | 0 | 0 | 0 |
| 36 | AF4 | 60 | 0 | 0 | 0 | 0 | 0 | 0 | 0 | 1 | 0 | 0 | 0 | 0 | 0 | 0 | 0 | 0 | 0 | 1 | 0 | 0 | 0 | 0 | 0 | 1 | 0 | 0 | 0 |
| 37 | wash | 30 | 1 | 0 | 0 | 0 | 0 | 0 | 0 | 0 | 0 | 0 | 0 | 0 | 0 | 0 | 0 | 0 | 0 | 1 | 0 | 0 | 0 | 0 | 0 | 1 | 0 | 0 | 0 |
| 38 | F1 | 60 | 0 | 0 | 1 | 0 | 0 | 0 | 0 | 0 | 0 | 0 | 0 | 0 | 0 | 0 | 0 | 0 | 0 | 1 | 0 | 0 | 0 | 0 | 0 | 1 | 0 | 0 | 0 |
| 39 | wash | 30 | 1 | 0 | 0 | 0 | 0 | 0 | 0 | 0 | 0 | 0 | 0 | 0 | 0 | 0 | 0 | 0 | 0 | 1 | 0 | 0 | 0 | 0 | 0 | 1 | 0 | 0 | 0 |
| 40 | AF3 | 60 | 0 | 0 | 0 | 0 | 0 | 1 | 0 | 0 | 0 | 0 | 0 | 0 | 0 | 0 | 0 | 0 | 0 | 1 | 0 | 0 | 0 | 0 | 0 | 1 | 0 | 0 | 0 |
| 41 | wash | 30 | 1 | 0 | 0 | 0 | 0 | 0 | 0 | 0 | 0 | 0 | 0 | 0 | 0 | 0 | 0 | 0 | 0 | 1 | 0 | 0 | 0 | 0 | 0 | 1 | 0 | 0 | 0 |
